# Genotype-phenotype mapping identifies fetal-like CD55^+^ immunoregulatory cancer cells as mediators of immune escape in colorectal cancer

**DOI:** 10.64898/2026.01.09.697991

**Authors:** Manuel Mastel, Aitana Guiseris Martinez, Gabriele Diamante, Jasmin Meier, Luzi Schuchmann, Nikolaos Georgakopoulos, Ioannis Chiotakakos, Laura K Steffens, Carolin Artmann, Diego Benitez, Michael Günther, Vera Thiel, Roxana Pincheira, Andreas Trumpp, Rienk Offringa, Steffen Ormanns, Rene Jackstadt

## Abstract

Immune evasion is a defining feature of advanced colorectal cancer (CRC), yet the cellular mechanisms linking cancer cell states to immune suppression remain poorly understood. To systematically map tumour-immune interactions in advanced CRC, we modelled recurrent human CRC mutations using a multiplex CRISPR-based genetically engineered mouse model platform. The resulting 20 models recapitulate key stages, genetic routes and histopathological features of human CRC, while single-cell transcriptomics reveals extensive disease complexity across models. Profiling of the epithelial tumour compartment identified a population of fetal-like immunoregulatory cancer cells (IRCs) characterized by interferon-γ and MAPK activity. IRCs are marked by expression of the membrane-bound complement regulatory protein CD55. Mechanistically, CD55 protects IRCs from immune surveillance and complement-mediated lysis. Together, these findings reveal a cancer cell-immune interface that promotes immune evasion in advanced CRC.

## Introduction

Colorectal cancer (CRC) is the second most prevalent cancer malignancy worldwide, with a well-defined mutational landscape^1,2^. While extensive genomic studies have identified critical driver mutations, their functional impact on tumour microenvironment remodelling remains poorly understood. Investigating how specific genetic alterations shape cancer cell states and modulate immune responses is critical for developing more effective therapeutic strategies^3^.

Immunotherapy has revolutionized the treatment of cancer, particularly in microsatellite instable CRC^4,5^. However, therapeutic strategies targeting immunoregulatory components for advanced microsatellite stable CRC are lacking. While individual driver mutations, such as *Kras*^6^, *Notch1*^7^ or *Smad4*^8^ have been reported to rewire the tumour immune microenvironment (TIME), a comprehensive view of how recurrent driver mutations collectively shape the tumour ecosystem is missing. Transcriptional subtyping identified four distinct consensus molecular subtypes (CMSs) of CRC^9^, in which the presence of lymphocytes is detected in CMS1 tumours and stromal cells defines the CMS4-subtype with the poorest clinical outcome. In contrast CMS2 and CMS3 tumours were defined as WNT-high, the “canonical” or “metabolic” subtypes, respectively. Together, these findings highlight that CRC subtypes emerge from complex interactions between cancer cells and tumour-microenvironment (TME) components, which strongly influence subtype identity and disease behaviour. While cell line panels capture some features *in vitro*, more advanced models are needed to faithfully reflect tumour-TME interplay *in vivo*^10^. In addition to the heterogeneity observed in non-cancerous cells within a tumour, cancer cells display a similarly complex composition^11,12^. Transcriptional subtyping on a single cell level revealed a WNT-high cell state, classified as intrinsic transcriptional subtype 2 (iCMS2), as well as cells characterized by MAPK-activation (iCMS3). Notably, iCMS3 is associated with the poorest prognosis^9,13^. The identity of cells along this continuum of cancer cell states was further characterized, revealing a trajectory from adult intestinal-like cells, marked by high WNT-activity and Lgr5-expression, to Yap- and MAPK-high, fetal-like cells associated with regenerative programs^14,15^. Importantly, cells in this state have been associated with drug resistance^16–20^ and metastatic recurrence^21^, positioning these cells as key contributors to CRC progression and potentially important therapeutic targets.

A major limitation in dissecting the heterogeneity of CRC is the lack of preclinical models that capture the genetic diversity of advanced tumours, their autochthonous progression from normal tissue to cancer, as well as their interactions with the TIME^22,23^. Traditional GEMMs provide insights into CRC initiation, but are slow to breed and inflexible when it comes to creating multiplexed genetic perturbations^7,24,25^. Patient-derived xenografts (PDXs) retain human tumour heterogeneity but lack a functional immune system^26^. Organoids derived from GEMMs represent a valuable addition to the current repertoire of models, particularly when transplanting in immunocompetent recipients^7,8,27^, but still fail to fully recapitulate the complexity of tumour progression from normal to pre-cancerous and cancerous stages. CRISPR-based *in vivo* genome engineering overcomes these limitations, enabling rapid, scalable, and combinatorial mutations in autochthonous settings to study tumour evolution from a normal cell to advanced cancer and immune evasion. Although applied successfully to generate benign lesions in the colon^28^ and in several other solid cancersc^29–36^, CRISPR/Cas9 strategies for advanced CRC have remained underdeveloped, despite the need to elucidate the mechanisms underlying the disease’s high heterogeneity, immune evasion, and complex microenvironment.

In this study, in addition to classical Cre/LoxP driven GEMMs, we developed a CRISPR-based *in vivo* genome engineering technology, designed to model advanced CRC directly *in vivo*. With this technology, we combined SOmatic CRISPR-mediated genome Alterations of Tumour suppressors with Explanatory single-cell Sequencing (SOCRATES), which enables rapid, combinatorial editing of key driver genes in colonic epithelium, bypassing the need for transgenic breeding. By integrating single-cell transcriptomics with *in vivo* genome editing, SOCRATES dissects complexity of cancer cell states and associated immune remodelling. Using this approach, we identify a fetal-like cancer cell population with high immunoregulatory capacity, defined by the complement regulatory receptor CD55.

## Results

### **Modelling the genetic landscape of advanced CRC** *in vivo*

To functionally model recurrent genetic drivers of advanced CRC^37^, we used two complementary approaches. First, we generated GEMMs that recapitulate the classical and serrated route to CRC driven by *Apc*, *Braf* or *Kras* mutations (Fig. 1a-b; Extended Data Fig. 1a). To induce tumour initiation, we used either systemic application of tamoxifen or colonoscopy-guided needle injection of 4-hydroxy tamoxifen (4-OHT). Second, we developed SOCRATES, a CRISPR-based technology for multiplexed *in vivo* genome engineering of advanced CRCs (Fig. 1a). SOCRATES utilizes a lentiviral system to simultaneously target *Apc* and *Trp53* while inducing *Kras*^G12D^ expression, generating combinatorial human-relevant driver mutations^37^. Additional sgRNAs targeting tumour suppressor (sgTS) or other genes of interest can be inserted, allowing for flexible genetic modifications (Extended Data Fig. 2a). This enables rapid and scalable introduction of gene of interest mutations, substantially accelerating classical Cre/LoxP-dependent transgenic breeding strategies.

**Figure 1.**
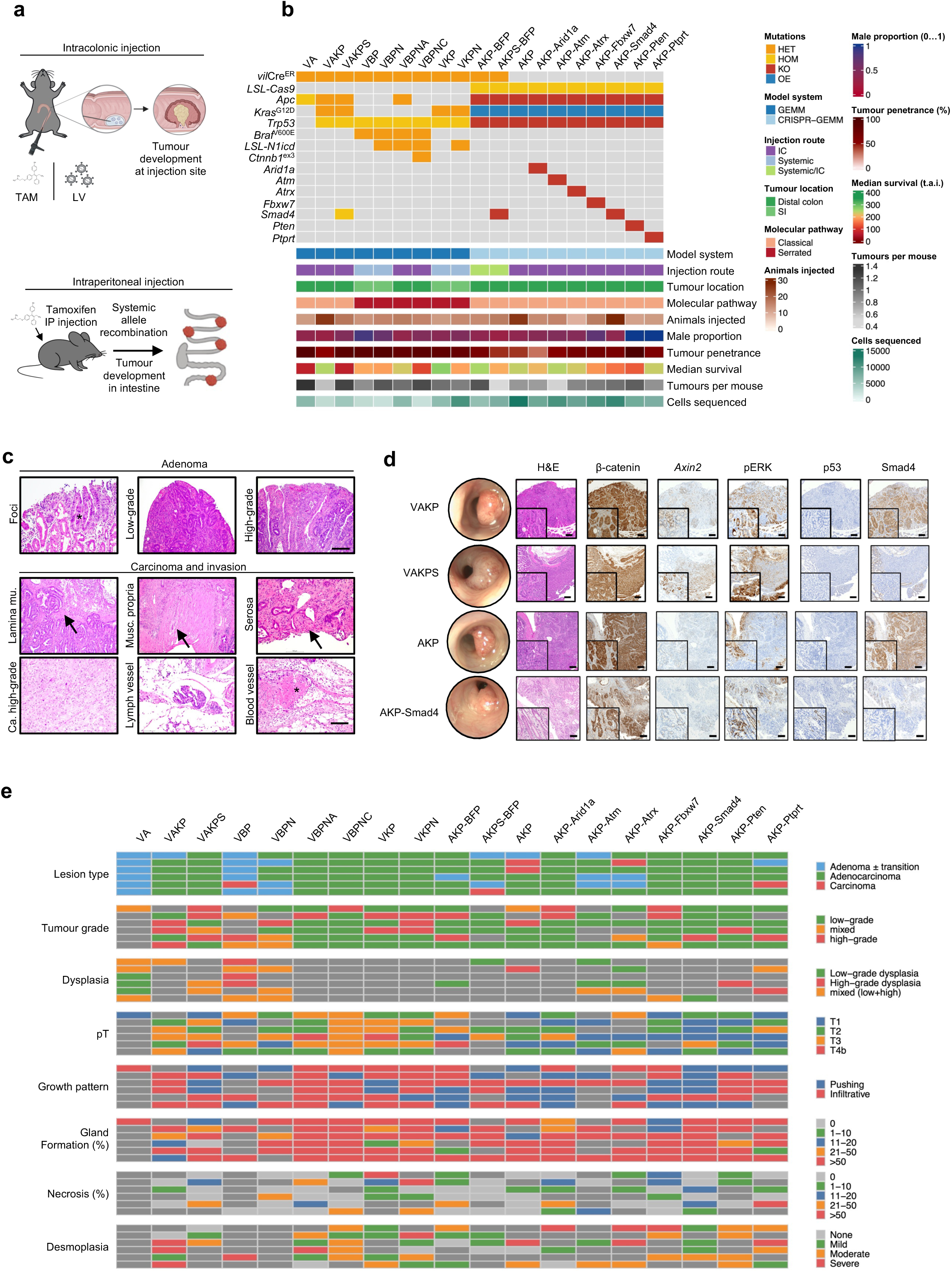
Comprehensive analysis of the genetic CRC mouse models. **a,** Schematic showing the induction routes to generate genetic compound mice. Intra peritoneal (IP); Tamoxifen (TAM); Lenti virus (LV). **B**, Specific features of genetic models of CRC. Knock out (KO); Over expression (OE); Intra colonic (IC); Small intestine (SI); Time after injection (t.a.i.). Exact values are provided in Extended Data Table 3. **C**, Histological appearance of models in early tumours with adenomatous foci (left), low-grade adenoma (middle) and high-grade adenoma (right). Scale bar 100 µm. Asterix indicates adenomatous foci. Carcinomas and invasion of SOCRATES-derived tumours. In order: Invasion to lamina muscularis, muscularis propria, serosa, high-grade carcinoma, lymph vessel and lymph blood vessel. Scale bar 100 µm. Arrows indicate invasion and asterisk indicates blood vessels. **D,** Representative colonoscopy pictures and biomarker immunohistochemistry of indicated genotypes. **E,** Quantification of histopathology of tumours across the genetic models.

To enable this technology, we generated a CRISPR-GEMM incorporating *Villin1*^CreER^ alongside the *Rosa26*^LoxP-Stop-LoxP-Cas^^9-GFP^ (LCas) transgene. Virus titre of this SOCRATES1 system was monitored with an integrated BFP reporter in CT26 cells (Extended Data Fig. 2a-b). To recombine the LCas allele, tamoxifen was administered systemically to the mice prior to local virus injection (Extended Data Fig. 2c). To test SOCRATES *in vivo*, we generated lentiviral particles encoding different CRISPR combinations and delivered them into the submucosa of *Villin1*Cre^ER^ LCas mice via colonoscopy-guided injections (Extended Data Fig. 2c-d).

To further increase versatility, we tested the efficacy of epithelial recombination in wild-type normal mucosa organoids derived from LCas mice. Organoids were infected with a Cre-recombinase driven by a PGK- or STAR-promoter which is composed of a minimal Lgr5-promoter element and 4x Ascl2 binding sites^38^. Both Cre-drivers led to robust recombination in organoids (Extended Data Fig. 3a-d). Hence, we replaced the BFP reporter cassette with STAR-Cre and generated an additional CT26 reporter system for virus titre analysis (Extended Data Fig. 3e-h). Analysis of indel frequency in 2D and 3D cell lines generated from tumours confirmed highly efficient genome editing using SOCRATES2, with robust indel formation across target loci (Extended Data Fig. 3i-m). Immunohistochemical analysis of biomarkers further validated the editing of targeted genes at protein level (Fig. 1d). GFP expression confirmed successful recombination, specifically in tumour cells but not surrounding normal mucosa (Extended Data Fig. 3k-n). Hence, SOCRATES2 with LCas mice represents strong advantages, including faster model generation and higher flexibility for modelling and studying advanced CRC.

Histopathological analysis of the generated tumour models confirmed stepwise progression from early adenomatous lesions to invasive carcinomas, with tumours invading deeper layers, including the muscularis propria and serosa (Fig. 1c-e). Two models *vil*^CreER^, *Apc*^fl/lf^ (VA) and *vil*^CreER^, *Brat*^V600E^, *Trp53* (VBP) were adenomas with low grade or high-grade dysplasia, respectively. While most of the other tumours resembled low-grade colorectal adenocarcinoma, a subset such as *vil*^CreER^, *Kras*^G12D^, *Trp53, Rosa26*^N1icd^, (VKPN) and *vil*^CreER^, *Brat*^V600E^, *Trp53, Rosa26*^N1icd^, *Ctnnb1*^ex^^3/+^ (VBPNC) showed high-grade features, including dysplasia. Key hallmarks of aggressive disease were also observed, such as lymphatic and vascular invasion, demonstrating that the model collection faithfully recapitulates the entire progression of CRC pathology *in vivo* (Fig. 1e). Notably, this approach generates a broad spectrum of CRC from low-grade adenoma to high-grade desmoplastic advanced CRC, closely mimicking human features and progression characteristics. Together, this genetically controlled model atlas recapitulates the heterogeneity and histopathology of CRC primary tumour progression and provides a robust foundation for systematic interrogation of tumour ecosystem complexity.

### Profiling of the tumour ecosystem in CRC models

To investigate how tumour suppressor alterations shape the tumour ecosystem, we performed single-cell sequencing on 131,716 cells derived from 126 samples including primary tumours across genotypes and normal mucosa (Fig. 2a-b). We recovered a heterogenous epithelial cell cluster and various clusters of non-malignant tumour cells such as T cells, endothelial cells, fibroblasts and myeloid cells (Fig. 2c; Extended Data Fig. 4a-e). Notably, we observed genotype-dependent shifts in cellular composition, with VA adenoma being enriched in epithelial cells, AKP-*Ptprt* tumours displaying increased stromal infiltration and AKP-*Atrx* tumours exhibiting a higher proportion of T/NK cell infiltration (Fig. 2d-e; Extended Data Fig. 5a). Overall, we found that tumours of the serrated route show more similarity compared to tumours of the classical route, indicating differences in the biology of these routes to CRC (Fig. 2e-f). Detailed annotation of cell states and polarizations within the TME uncovered a high degree of cellular complexity comparable to that observed in human CRCs (Extended Data Fig. 4e)^39^. To define the CMS subtype of the respective models we set out to apply the CMS classification on our cohort of single-cell data. However, utilizing available tools for mouse bulk datasets resulted in a large proportion of unassigned tumours (Extended Data Fig. 5b-g). To circumvent this, we curated balanced CMS-like gene sets in mouse orthologues capturing immune/MSI (CMS1), canonical WNT/MYC–epithelial (CMS2), metabolic (CMS3), and mesenchymal/stromal-TGFβ programs (CMS4) (Extended Data Table 1) and we computed per-sample enrichment scores for each CMS program by single-sample GSEA (GSVA/ssGSEA). We then assigned each sample a discrete “CMS-like” call by selecting the program with the highest ssGSEA score. This analysis revealed that the majority of VA adenomas were assigned to the WNT-high subtypes CMS2/3, aligning with high WNT-pathway activity of adenoma (Extended Data Fig. 5h). Interestingly, the SOCRATES2 model AKP-*Pten* showed the strongest CMS4 phenotype. Overall, these data demonstrate that genetic alterations exert a dominant influence on the tumour ecosystems composition and that our dataset captures key features of CRC heterogeneity.

**Figure 2.**
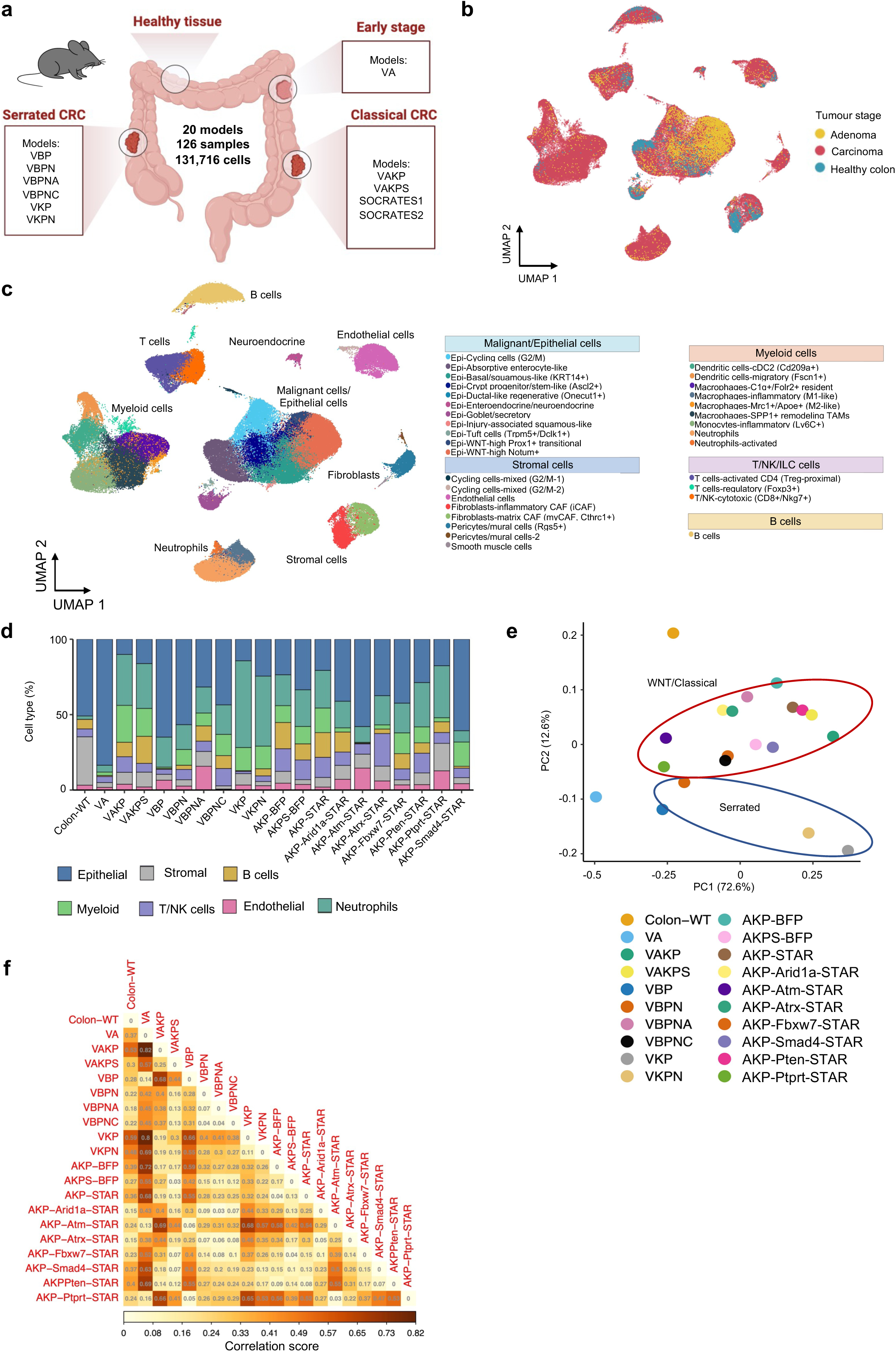
Transcriptional profiling of the genetic mouse models. **a,** Cartoon indicating the models of the atlas that were analysed by single cell transcriptomics. **b,** UMAP plot showing cellular contribution across stage. **c,** UMAP plot showing cell type annotation (131,716 cells; 126 samples). **d,** Proportions of cell types of the atlas derived tumours with indicated genotypes. **e,** PCA of tumour models from d. **f,** Heat-map showing the distances between each two tissue types.

### Characterisation of immunoregulatory cancer cells in CRC

Next, we sought to chart cancer cell heterogeneity across the models. To this end we focused on cells expressing epithelial marker genes that showed a clear distinction from other cell types. Unsupervised clustering uncovered 14 distinct clusters (Fig. 3a). Most recognisable were cells with neuroendocrine differentiation (marked by *Neurod1*), ductal-like regenerative features (marked by *Onecut1*) and secretory linages such as goblet like (marked by *Muc2*) and absorptive enterocyte-like cells (marked by *Fabp1*) (Fig. 3a; Extended Data Fig. 6a). Normal tissue and adenoma were assigned to goblet-like cells and WNT-signalling high populations (Fig. 3b), confirming previously detected strong WNT-activity in adenomas^40,41^. Epithelial cell intrinsic subtyping revealed that cells in the WNT and cell cycle cluster belong to iCMS2, while stress and injury cluster cells were assigned to iCMS3 (Fig. 3c), recapitulating the distribution in human CRC^13^. Consistent with CMS-like- assignments, AKP-*Pten* showed the strongest iCMS3 assignment and VA adenoma the most robust iCMS2 association, thus validating subtyping strategies (Fig. 3c; Extended Data Fig. 6b-c). Intriguingly, genotype-specific differences in epithelial cell-state composition were observed, indicating that oncogenic drivers actively shape cell-state composition (Fig. 3d; Extended Data Fig. 6d). Tumours arising through the serrated route, including VBPN, VKP and VKPN, exhibited highly similar cell-state distributions, with increased proportions of squamous-like epithelial cells. Adenomas (VA) appeared most strikingly different to the serrated tumours in a principal component distribution analysis (Fig. 3e). This analysis revealed a population, termed immune regulatory epithelium, that showed an enrichment for stress- and immune-signatures at the same time (Fig. 3f). Further characterization of these cells showed activated programs of fetal-like, Yap-high and regenerative-signatures, but at the same time association to immune response signatures such as INFγ- activation and negative immune regulation (Fig. 3f-g; Extended Data Fig. 6e). Based on these combined features, we termed this cluster immunoregulatory cancer cells (IRCs). When we performed pseudo time trajectory analysis to define the progression trajectories of cancer cells, with a set starting point in normal tissue and intermediate in the adenoma stage, we identified four trajectories (Fig. 3h). These trajectories had their terminal state in distinct clusters of EMT, fetal, proliferation or IRCs (Fig. 3i). Tumours were stratified according to the proportion of cells occupying each terminal cluster (EMT, fetal-like, proliferative, or IRCs), and overall survival was compared between mice bearing tumours with high versus low cluster occupancy (Fig. 3j). Across both GEMM and CRISPR-GEMM cohorts, IRCs were associated with significantly reduced survival (Fig. 3j). Collectively, the complexity of the murine atlas enabled us to define previously described cell states and enabled the discovery of the IRC cluster.

**Figure 3.**
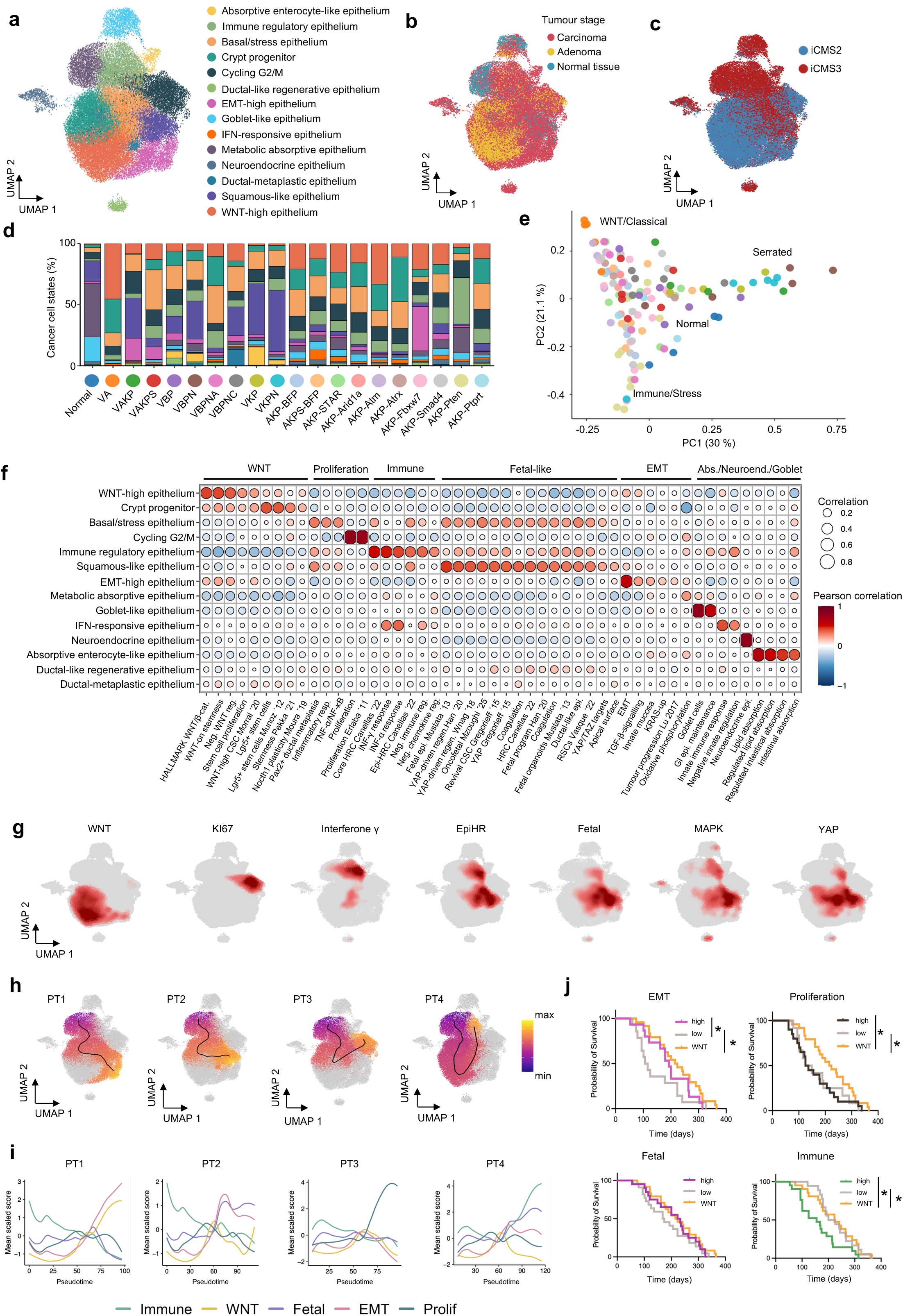
Epithelial cell states across stages and genotypes. **a,** UMAP plot showing epithelial cell subsets (n=44,594 cells). **b,** UMAP plot showing tumour stages. **c,** UMAP plot showing tumour stages with iCMS subtyping **d,** Bar graph showing cellular distributions across models. **e,** PCA plots showing sample clustering based on cell subset abundance, coloured by model. **f,** Bubble plot showing gene-set enrichment across epithelial clusters. **g,** UMAP plots showing pathway activity in epithelial cells. **h,** Pseudo time (PT) trajectory in epithelial clusters. **i,** Trajectory analysis for indicated pathways/processes. **j,** Kaplan-Meier plots of the indicate pathways/process as identified in h. Survival was compared using a log-rank (Mantel-Cox) test. Two-sided *p*-values are shown.

### Tumour immune infiltrate in IRC tumours

Given the identification of IRCs, we aimed to understand their impact on the tumour-ecosystem. To gain insight into IRCs identity in the cancer cell compartment, we performed gene-set enrichment analysis of IRCs compared to non-IRC epithelial cells. This revealed strong enrichment for interferon signalling, immune regulatory pathways, and epithelial stress responses (Fig. 4a-b). The serrated models VKPN and VBPN, as well as normal colon and adenoma (VA), exhibited the lowest abundance of IRCs, whereas AKP-*Pten*, VBPNA, and AKPS displayed the highest proportion of IRCs within the cancer cell compartment (Fig. 4c). To assess the association of IRCs with the tumour immune infiltrate, we compared its composition between tumours with high- and low-IRC content. While the overall composition of cell types didn’t change remarkably, we detected a heterogenous polarization of myeloid cells and a significantly smaller fraction of reactive macrophages in IRC-high tumours (Fig. 4d-e). Interestingly, similarly to the IRC epithelial cluster, macrophages showed an increase in INFγ-signatures, indicating more INFγ-activation in IRC-high tumours. In addition, we observed an enrichment of innate immune response pathways in macrophages of IRC-high tumours, whereas IRC-low tumours displayed enriched macrophage activation and inflammatory responses (Fig. 4f). Moreover, neutrophils in IRC-high tumours exhibited a pro-tumorigenic myeloid-derived suppressor cell (MDSC)-like state, accompanied by reduced inflammatory, complement and immune responses activation in IRC-low tumours (Fig. 4g-i). Together, these data indicate a chronically inflamed tumour microenvironment under active immune pressure in IRC-high tumours that is counterbalanced by robust neutrophil-mediated immunosuppression.

**Figure 4.**
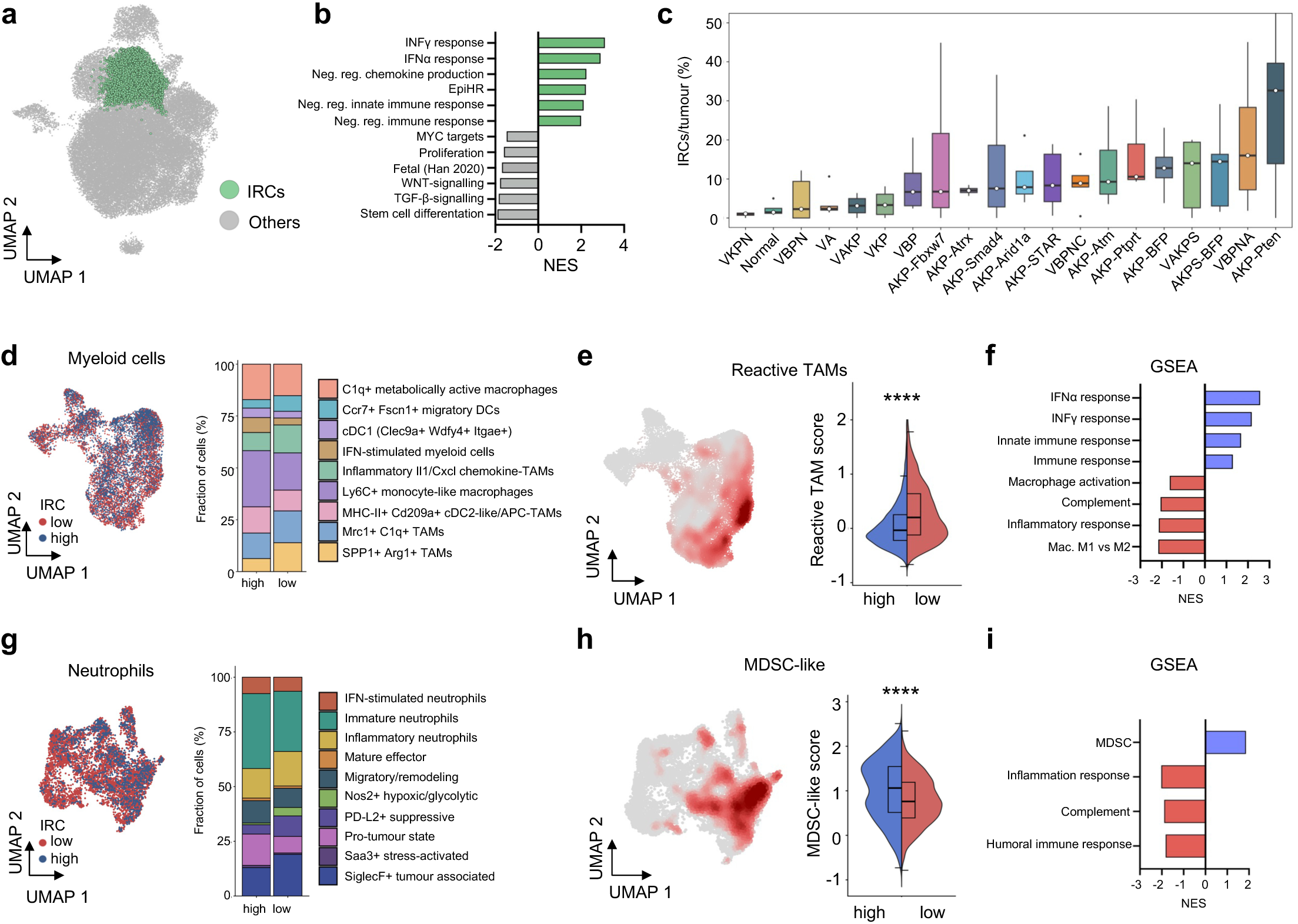
IRC tumour ecosystems. **a,** UMAP plot highlighting the immune regulatory cancer cell (IRC) population in the epithelial cell subset (n=4,004 cells). **b**, Gene-set enrichment analysis of the IRC population compared to all other epithelial cells. **c**, Percentage of IRCs per tumour across the models. **d**, UMAP plot of re-clustered myeloid cells (left) and corresponding compositional analysis (right) classified by the IRC expression of the tumour of origin (n=32,285 cells). **e,** UMAP plot of re-clustered myeloid cells highlighting reactive TAMs and split violin box-plot comparing the score of reactive TAMs, between the IRC expression on tumour of origin. Statistical significance was assessed using a two-sided Wilcoxon Rank-Sum test. **F,** Gene-set enrichment analysis of myeloid cells from IRC-high tumours compared to IRC-low. **G**, UMAP plot of re-clustered neutrophils (left) and corresponding compositional analysis (right) classified by the IRC expression of the tumour of origin (n=11,616 cells). **h**, UMAP plot of re-clustered neutrophils highlighting MDSC-like neutrophils and split violin box-plot comparing the score of MDSC-like neutrophils, between the IRC expression on tumour of origin. Statistical significance was assessed using a two-sided Wilcoxon Rank-Sum test. **I**, Gene-set enrichment analysis of neutrophils from IRC-high tumours compared to IRC-low.

### CD55 defines IRCs with complement inhibitory capacity

To explore the role of IRCs in immune regulation, we sought a candidate marker to distinguish this population. As aforementioned we detected signatures of immune enterocytes marked by *Lgals4* or *Cecam1*, complement regulation marked by *Cd55* and interferon or antiviral interferon signalling genes marked by *Irf7* (Fig. 3f; Extended Data Table 1). Among these candidates, the membrane-bound complement regulatory protein CD55 showed the strongest enrichment and correlation with the IRC transcriptional signature (Fig. 5a-c). We tested the expression of *Cd55* across the tumour models of the atlas and found high expression in VBPNA, VAKPS and VBPNC tumours and low expression in VKP or VBPN (Fig. 5d). We confirmed surface expression of CD55 in organoids with respective genotypes (Fig. 5e). To test the regulation of *Cd55,* we treated low expressing VBPN organoids with EGF or INFγ and found a time dependent induction with both compounds (Fig. 5f). Similarly, canonical MAPK target genes, such as *Egr1*, and IFNγ-responsive genes, such as *Cd274*, were also induced (Fig. 5g).

**Figure 5.**
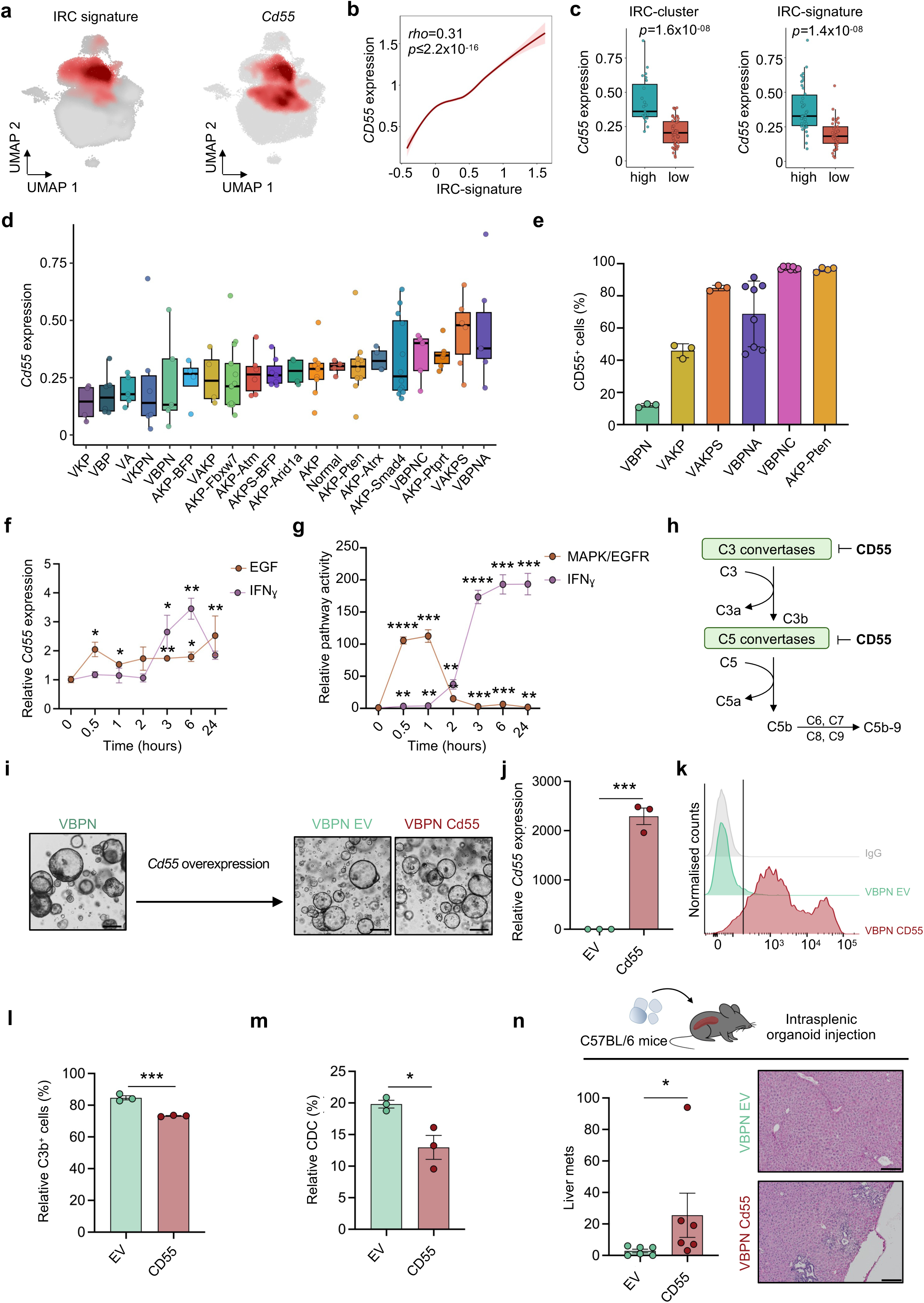
CD55 marks IRCs with reduced complement immune surveillance. **a,** UMAP plot showing the comparison of immune regulatory cells (IRCs) and *Cd55* expression in the epithelial cell subset. **b**, Correlation plot between IRCs and *Cd55* expression. Spearman’s rank correlation coefficient (ρ) and two-sided *p*-values are indicated. Linear regression or LOESS smoothing (all epithelial cells) is shown for visualization only. **c**, *Cd55* expression in tumours containing a high or low proportion of cancer cells in the IRC-cluster. *Cd55* expression of tumour cells containing a high or low IRC-signature expression. **d**, *Cd55* expression across the epithelial cell subset of the models. Differences across models were assessed using a Kruskal-Wallis test; the two-sided p-value is shown. **e,** Flow cytometric quantification of CD55 expression across organoids derived from different models (n=3, 3, 3, 8, 8, 4). **f,** qPCR analysis of VBPN cells treated with the indicated cytokine (n=3). **g,** qPCR analysis of VBPN cells treated with indicated reagents as in f for *Egr1* (MAPK) and *Cd274* (INFγ) (n=3). **h**, Schematic illustrating the role of CD55 in regulating the complement cascade. **i**, Schematic of the lentiviral overexpression strategy and representative images of the resulting organoids. Scale bar 250 µm. **j**, qPCR analysis of *Cd55* expression in VBPN EV and VBPN CD55 cells (n=3). **k**, Histogram showing *Cd55* expression in VBPN EV and VBPN CD55 cells. **l,** Cell-surface C3b deposition relative to heat-inactivated serum (HIS), expressed as a percentage (n=3). **m**, Relative complement-dependent cytotoxicity (CDC) normalized to HIS (n=3). **n**, Number of metastases in C57BL/6 mice (n=6) and representative haematoxylin and eosin-stained liver sections. Scale bar 200 µm. Statistical significance was assessed using a Mann-Whitney test. Statistical significance in c, f, g, j, l and n was assessed using an unpaired Student’s *t*-test.

Next, we aimed to investigate the functional role of CD55 in CRC progression and its potential negative regulation of the complement cascade through acceleration of C3 and C5 convertase decay (Fig. 5h), which can promote pro-tumorigenic phenotype via complement inhibition. Hence, we transduced the low CD55-expressing VBPN organoid line with lentiviruses encoding for the murine *Cd55* ORF (VBPN CD55) or the corresponding empty vector (VBPN EV) (Fig. 5i-k; Extended Data Fig. 7a-b). Consistently with this hypothesis, CD55 overexpression attenuated complement activation, as reflected by reduced C3b deposition (Fig. 5l) and significantly decreased complement-dependent cytotoxicity (CDC) (Fig. 5m). Intra-splenic transplantation of VBPN EV or VBPN CD55 organoids into immunocompetent mice resulted in an increased metastatic burden (Fig. 5n). Together, this data demonstrates induction of CD55 by INFγ and EGF and shows that CD55 promotes metastatic dissemination.

### CD55-mediated tumour immune remodelling

To understand the role of CD55 in shaping the TIME, we next deleted CD55 in two organoid lines exhibiting high CD55 protein expression (Fig. 5e; Fig. 6a-b; Extended Data Fig. 7c-d). Transplantation of isogenic organoids with CRISPR/Cas9-mediated *C055* knock out (KO) or non-targeting controls (NT) led to reduced tumour growth and metastatic seeding in immunocompetent mice (Fig. 6c-d; Extended Data Fig. 7e-f). To test whether this effect was complement-dependent, we performed this assay in *C3*-deficient mice. Mice lacking C3 are unable to produce anaphylatoxins that may lead to infiltration of immunoregulatory myeloid cells. Indeed, the effect of *Cd55* depletion was rescued in *C3*^-/-^ mice (Fig. 6e). To examine whether the deletion of *Cd55* led to <u>distinct</u> immune cell infiltration, we performed Cellular Indexing of Transcriptomes and Epitopes by sequencing (CITE-seq) on tumours from C57/BL6 mice. The immune CITE-seq panel, comprising a cocktail of 102 oligonucleotide-conjugated antibodies, included an antibody detecting CD55, which confirmed loss of CD55 at the protein level, consistent with reduced *Cd55* transcript abundance (Extended Data Fig. 7g). Profiling recovered the major tumour-associated cell populations, including myeloid cells, T cells, fibroblasts, and endothelial cells (Fig. 6f). Notably, *Cd55* knockout tumours displayed marked increase in macrophage abundance (Fig. 6g-h). Further phenotypic characterisation detected an increase of the anaphylatoxin receptor C5aR1⁺ expressed on macrophage and neutrophil in *Cd55* knockout tumours. These populations dominantly exhibiting anti-tumour phenotypes, such as inflammatory neutrophils (Fig. 6i-l). Collectively, these data indicate that CD55 is not only sufficient but also required for tumour progression, and that CD55 shapes the infiltration and phenotype of immune cells via complement inhibition.

**Figure 6.**
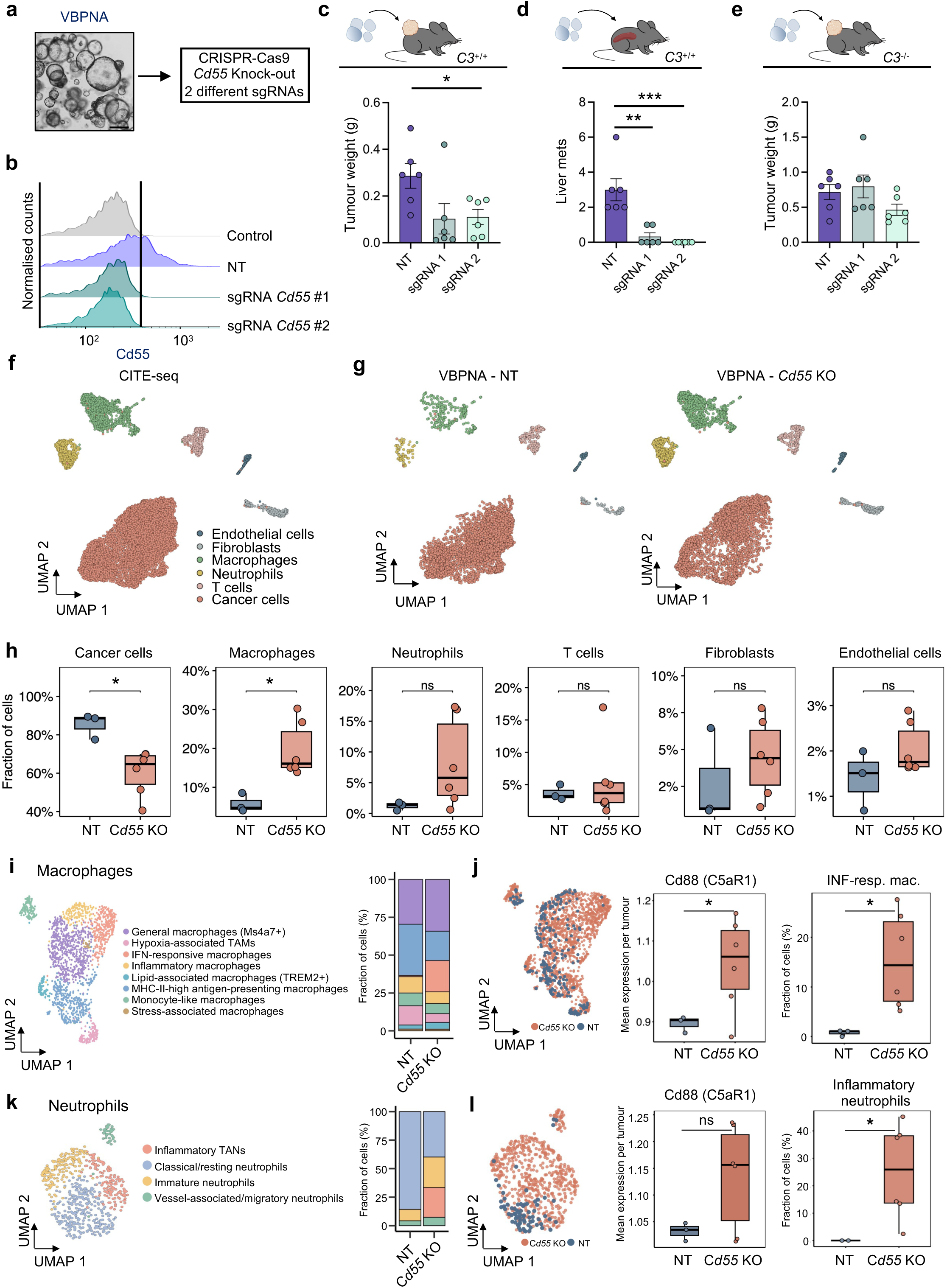
CD55 controlled TIME rewiring. **a,** Schematic describing the CRISPR/Cas9 knock-out strategy. **b,** Histogram of *Cd55* expression in non-targeting sgRNA or *Cd55* sgRNAs. **c,** Tumour weights at endpoint of transplanted organoids from a in C57BL/6 mice (n=6). **d,** Number of metastasis in C57BL/6 mice (n=6). **e,** Tumour weights at endpoint of transplanted organoids from a in *C3*^-/-^ mice (n=6). **f,** UMAP plot showing CITE-seq analysis from c (n=13,549 cells). **g,** UMAP plot showing CITE-seq analysis non-targeting (NT) control versus VBPNA tumours and *Cd55* knock-out (KO). **h,** Box-plots showing the quantification of cell type distribution for NT (n=6) and *Cd55* KO (n=6). **i,** UMAP plot and quantification of macrophage polarization. **j,** UMAP plot and quantification of macrophage states. **k,** UMAP plot and quantification of neutrophil polarization. **l,** UMAP plot and quantification of neutrophil states. Statistical significance in c, d, e, h, j and l was assessed using a two-sided unpaired Student’s *t*-test.

### IRCs and clinical features in human CRC

To validate the mouse CRC atlas in human CRC samples we used a dataset of 459,557 cells comprising 312 samples (Fig. 7a)^39^. Similar to the murine atlas we extracted the data of normal tissue, adenoma and carcinoma (Fig. 7b). The cell type composition was similar to the murine atlas and included epithelial cells, endothelial cells, fibroblasts and various immune cell clusters (Fig. 7c). Site-by-site comparisons revealed concordance between mouse and human adenoma and carcinoma cellular compositions (Fig. 7d-e). Using a joint principal component analysis (PCA) based exclusively on tumour cell-type composition, in which samples were represented by relative abundances of annotated cell populations, we observed high degree of similarity between human and mouse tumours across disease stages (Fig. 7f). We found that CD55 expression strongly overlapped with the IRC signature derived from the murine atlas (Fig. 7g-h). Expression of CD55 was increased in carcinomas, compared to normal tissues and adenomas (Fig. 7i). Using TCGA-COAD bulk RNA-seq data, we aggregated IRC-marker genes into an IRC-derived bulk expression signature (Extended Data Table 1) and found that elevated IRC-signature expression was associated with reduced overall survival (Fig. 7j). Consistent with immune- and stromal-rich subtypes, the IRC-signature was enriched in CMS1 and CMS4 tumours (Fig. 7k). Moreover, IRC-gene signature scores increased with advancing American Joint Committee on Cancer (AJCC) pathologic stage, showing a monotonic association with disease progression from stage I to IV (Fig. 7l). Together, these analyses demonstrate that IRCs represent a conserved, CD55-associated tumour cell state in human CRC that mirrors the murine atlas, are linked to adverse clinical features, and are associated with immune-enriched tumour subtypes, advanced disease stage, and poor patient outcome.

**Figure 7.**
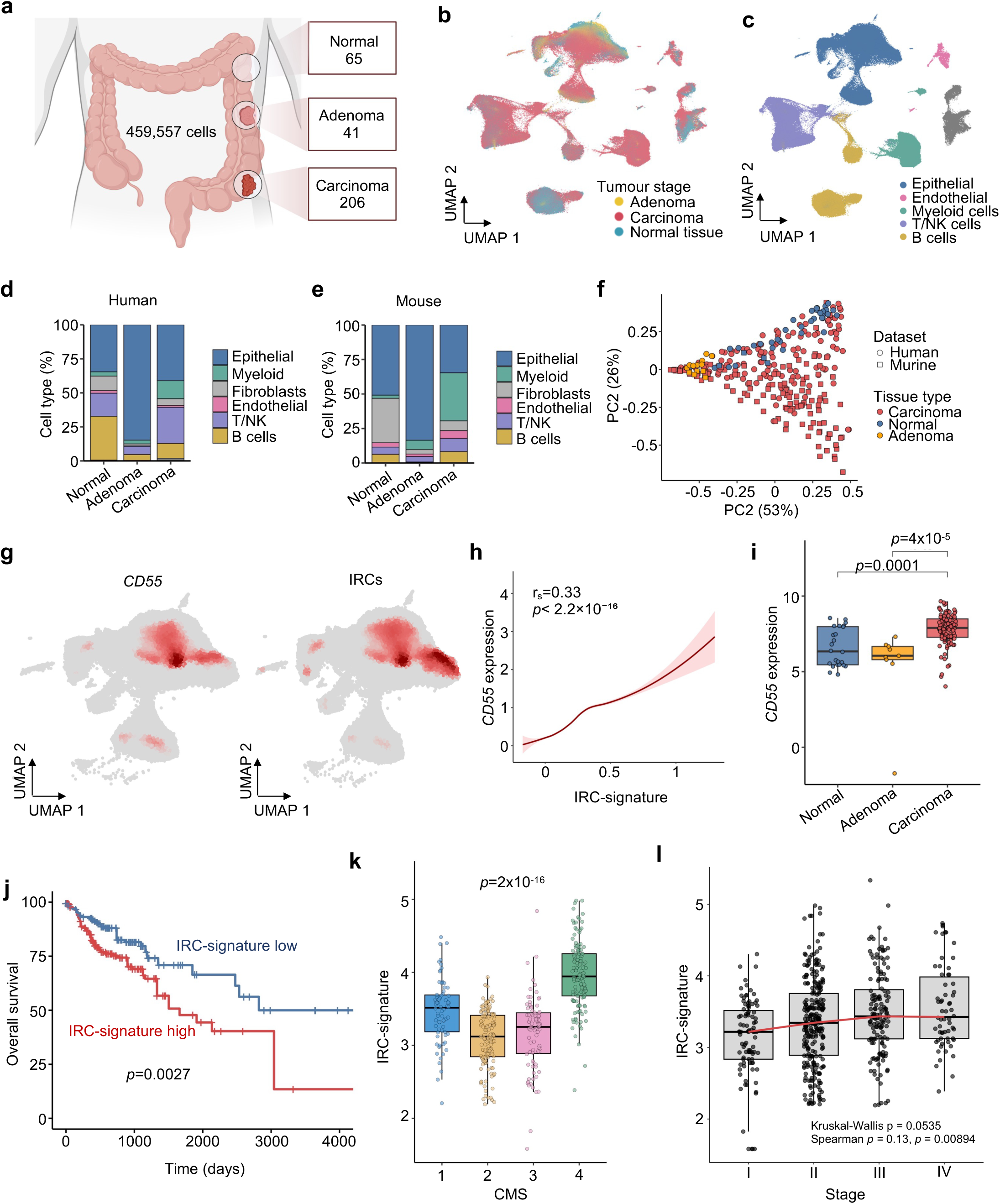
Characterization of IRCs in human CRC. **a,** Cartoon describing the human single cell transcriptomics cohort. **b,** UMAP plot showing the cell type distribution across stages (n=459,557 cells). **c,** UMAP plot showing cell type annotations. **d,** Bar plot showing cell type distribution across stages in human CRC cohorts (65 normal tissue, 41 adenoma and 206 carcinoma samples). **e,** Bar plot showing cell type distribution across stages in the mouse CRC atlas. **f,** PCA plots showing sample clustering based on cell subset abundance, coded by stage and species (n=438). **g,** UMAP plots showing *C055* and IRC expression in the epithelial subset of human CRC. **h,** Correlation of *C055* and IRC cluster expression in epithelial human CRC cells. Spearman’s rank correlation coefficient (ρ) and two-sided *p*-value is indicated. Linear regression (CD55⁺ cells) or LOESS smoothing (all epithelial cells) is shown for visualization only. **i,** Box-plot showing the expression of CD55 across stages in human CRC (normal n=65; adenoma n=41; carcinoma n=206). Differences across groups were assessed using a pairwise rank-sum test. Two-sided *p*-values are shown. **j**, Kaplan-Meier curve of IRC signature high and low tumour patients within the human COAD TCGA dataset with 470 patients. Survival was compared using a log-rank (Mantel-Cox) test. Two-sided *p*-value is shown. **k**, Boxplots, showing association of IRC signature score with CMS in the COAD TCGA dataset of 470 patients. Differences across CMS groups were assessed using a Kruskal-Wallis test; the two-sided *p*-value is shown. **l,** Boxplots showing association of IRC signature score increase with advancing AJCC stage in CRC. Overall differences between stages were assessed using a Kruskal-Wallis test and monotonic association with disease stage was evaluated using Spearman rank correlation. Exact *p*-values and correlation coefficients are indicated in the plot.

## Discussion

Understanding the biological complexity of advanced cancers requires a comprehensive view of how genetic alterations shape tumour progression and the interactions with the surrounding microenvironment. While mutations are key drivers of tumour initiation, their functional impact on tumour progression, particularly their influence on immune and stromal components and their ability to reshape the tumour ecosystem remains unclear^42,43^. Dissecting this landscape of driver mutations in advanced CRC is currently limited by the lack of advanced CRC mouse models. A comprehensive systems analysis using established and novel GEMMs, and the development of multiplexed CRISPR-GEMMs enabled us to functionally and mechanistically investigate the cellular and molecular determinants of CRC progression. Notably, tumours generated with either SOCRATES1, SOCRATES2 or conventional GEMMs showed concordance, demonstrating that the CRISPR-GEMMs recapitulate the Cre/loxP driven GEMMs. We utilized this array of models to get an unprecedented view on epithelial cell heterogeneity and identified a fetal-like immunoregulatory cancer cell population. IRCs influence immune surveillance of the innate immune system via the complement regulatory protein CD55. These mechanistic insights serve as an example of how powerful this collection of models is to identify and functionally dissect critical interactions within the tumour ecosystem in immune-competent models.

By applying single-cell transcriptomics to tumours from our murine atlas, we observed that driver mutations shape cancer cell states, immune cell polarization, and stromal remodelling. Notably, tumours following the classical and serrated routes of CRC exhibited the largest divergence in their ecosystem composition. These findings complement those of Househam *et al.*^44^, who reported that genetic variation only partially dictates gene expression programs in human CRC, with transcriptional plasticity contributing substantially to intratumour heterogeneity. In our controlled mouse models, the impact of defined driver mutations on cellular ecosystems can be directly observed, providing a mechanistic framework that supports and extends the human observations: while genetic background sets broad constraints on tumour evolution, the resulting transcriptional and microenvironmental states exhibit considerable plasticity.

Our subtyping analysis recovered CMS-like heterogeneity, with WNT-high tumours enriched for CMS2/3 signatures and serrated tumours for CMS1/4 signatures across driver combinations, in line with previous observations^7,45,46^. Interestingly, our collection reveals heterogeneous tumour-ecosystems across driver combinations, recapitulating the immunosuppressive tumour microenvironment observed in the majority of human CRC^8,9^. Similarly, to human CRC we found distinct states of CAFs, endothelial cells, myeloid cells, T cells, B cells and cancer cells^39^. Together, these findings demonstrate that genetic alterations orchestrate diverse immunosuppressive programmes.

Analysing the cancer cell compartment of the murine atlas showed that adenomas were defined by high WNT-signalling activity, while disease progression into advanced CRC genotypes exhibited a shift towards an increase of fetal-like regenerative programs. This observation is in line with previous observations in human CRC where, for example, TCF and LEF motifs had increased activity in polyps^40,41^ and progressive disease is characterised by enhanced cellular plasticity and fetal-like cell states^47^. Deeper characterization of the epithelial compartment allowed us to uncover distinct clusters such as EMT, WNT-high and various WNT-low clusters. Since WNT-low clusters were recently associated to the fetal-like regenerative programs in CRC, we investigated these states closer^48^. We became interested in IRCs since they showed simultaneously fetal-like, regenerative or YAP-high cell identity and exhibited INFγ-response signatures as well as immune evasion marker expression. INFγ mediated induction of fetal-like signatures was previously recognized during helminth infection, which induced loss of Lgr5 and stem cell remodelling in the small intestine^49^. In early tumorigenesis of mouse models, SOX17 produces LGR5-negative cells to evade anti-tumour immunity^50^. Our data is in line with this observation and shows expression of SOX17 in the adenoma clusters. Interestingly, recent clinical and pre-clinical biomarker analyses position IFNγ-responsive tumours, or those with increased IFNγ-signatures, as critical determinants for response to immune checkpoint inhibition^51,52^. However, our data indicates a complex immune infiltration in IRC-high tumours with strong IFNγ-response signatures. Hence, the role of IRCs in inflammatory responses and their potential relevance for immunotherapies warrant further investigation.

As part of the innate immune response, the complement system represents the first line of defence against pathogens. Thus, all human cells express surface molecules that regulate the activation of C3^53^. C3 convertase cleaves C3 into C3a and C3b, a rate-limiting step inhibited by CD55. Our data show that CD55 inhibits opsonin-promoting phagocytosis, and loss of CD55 enhances the recruitment of C5aR1-positive inflammatory myeloid cells. Given this central role of CD55 in the complement cascade, targeting CD55 may trigger context-dependent responses. In humanised mouse models of lung cancer, CD55 blockage in combination with PD-1 inhibition showed enhanced tumour suppressive efficacy^54^. Given the enhanced INFγ-activation in IRCs and the induction of CD55 and PDL-1 by INFγ in our organoids, these findings suggest that similar strategies of co-targeting CD55 and the PD-L1/PD-1 axis might overcome refractoriness to PD-L1/PD-1 monotherapies in CRC. Our observation that the complement regulatory protein CD55 marks IRCs suggests a central role for this cell type in CRC immunosuppression and the control of host immunity. The regulation and function of the complement system in cancer is context-dependent^53^. However, CD55 expression is associated with poor prognosis in CRC and our findings are in line with previous observations of enhanced CD55 expression in tumour tissues compared to surrounding normal tissues^55,56^. In addition to our observation that CD55 is induced by EGF and IFN-γ, CD55 has also been reported to be upregulated by prostaglandin E2 (PGE2) in CRC cell lines^57^. Interestingly, fibroblast-derived PGE_2_ triggers, via a Ptger4-Yap cascade, the appearance of Ly6a^+^ revival/fetal-like cells in tumour initiation^58^, suggesting that PGE_2_ might be an additional driver of the IRC state.

These findings indicate the importance of targeting particular cell-states and tumour phenotypes in advanced CRC. The flexibility to systematically dissect these states instructed by genotypes, in an autochthonous setting, represents a key advantage of our approach. While organoid generation and subsequent transplantation is a widely used and applicable strategy to generate immune-competent tumours, with strong assets over conventional 2D mouse derived CRC lines, it comes with a number of shortcomings. For example, only advanced stages of CRC can be modelled since orthotopic transplantation of precursor derived organoids is inefficient. Moreover, the progression from normal mucosa towards advanced CRC is not possible using organoid transplantation models.

Beyond its immediate applications, SOCRATES has the potential to be adapted to other epithelial cancers, for example sarcoma, gastric-, oesophageal-, skin-, and ovarian-cancers^59–65^. Future technological advancements could include base- or prime-editing for precise mutational design^34,66^, CRISPR barcoding to trace clonal tumour evolution at single-cell resolution and integration with patient-specific genetic variants^36,67^. Our CRISPR-based modelling strategy enables robust and scalable interrogation of cancer driver function *in vivo*; however, genetic perturbations may introduce heterogeneity due to variable editing efficiencies or rare unintended genomic alterations. Addressing these effects through orthogonal validation approaches, including independent sgRNAs and complementary genetic models, will further strengthen causal genotype-phenotype assignments and enhance mechanistic resolution in future studies.

Together, our technologies provide a versatile and scalable system for mapping genotype-phenotype relationships in solid tumours. This enables for high-resolution analysis of cancer evolution, immune interactions and potentially therapy resistance. Our atlas uniquely integrates somatic genome engineering with single-cell transcriptomics in an autochthonous setting. Compared to existing *in vivo* models, SOCRATES uniquely combines rapid genetic manipulation, complex tumour ecosystem modelling and the ability to track cancer progression. Our findings highlight that tumour progression is not solely dictated by intrinsic mutations, but by an interplay between genetic alterations, cancer cell state, and the surrounding ecosystem. Understanding these interactions will be critical for improving patient stratification and identifying novel therapeutic vulnerabilities in advanced cancers.

## Author contributions

M.M. designed the research, performed the research, analysed data and edited the manuscript. A.G.M. designed the research, analysed data, performed the research and edited the manuscript. G.D. performed the research. J.M. performed the research. L.S. performed the research. N.G. performed the research. I.C. performed the research. L.K.S. performed the research. C.A. performed the research. D.B. performed the research. M.G. performed the research. V.T. performed the research. R.P. provided resources. A.T. provided resources. R.O. analysed the data. S.O. performed the research and analysed the data. R.J. designed the research, performed the research, analysed data, wrote the first version of the manuscript, supervised the work and acquired funding. All authors read and approved the manuscript.

## Acknowledgements

The authors thank all Jackstadt laboratory members for discussions, the core units at the German Cancer Research Center (DKFZ), the animal facilities, transgenics services, sequencing facility and the flow cytometry facility. We thank Hugo J. Snippert for providing the STAR plasmid. Cartoons were created with BioRender. The work was supported by the Dietmar Hopp foundation and by the Dr. Rolf M. Schwiete Foundation (2021-032), the Deutsche Forschungsgemeinschaft DFG (JA 2558/3-1; JA 2558/4-1; JA 2558/6-1; JA 2558/9-1 P06 (FOR 5806)) and BMBF funded SATURN^3^ project (01KD2206B; 01KD2206E) to R.J. and the Joachim-Herz fellowship funded to M.M., T.T. and L.K.S.. Fellowships were provided by the Helmholtz International PhD Graduate School to M.M., G.D. and by the Merck’sche Society for Art and Science to A.G.M..

---

**Extended Data Figure 1.**
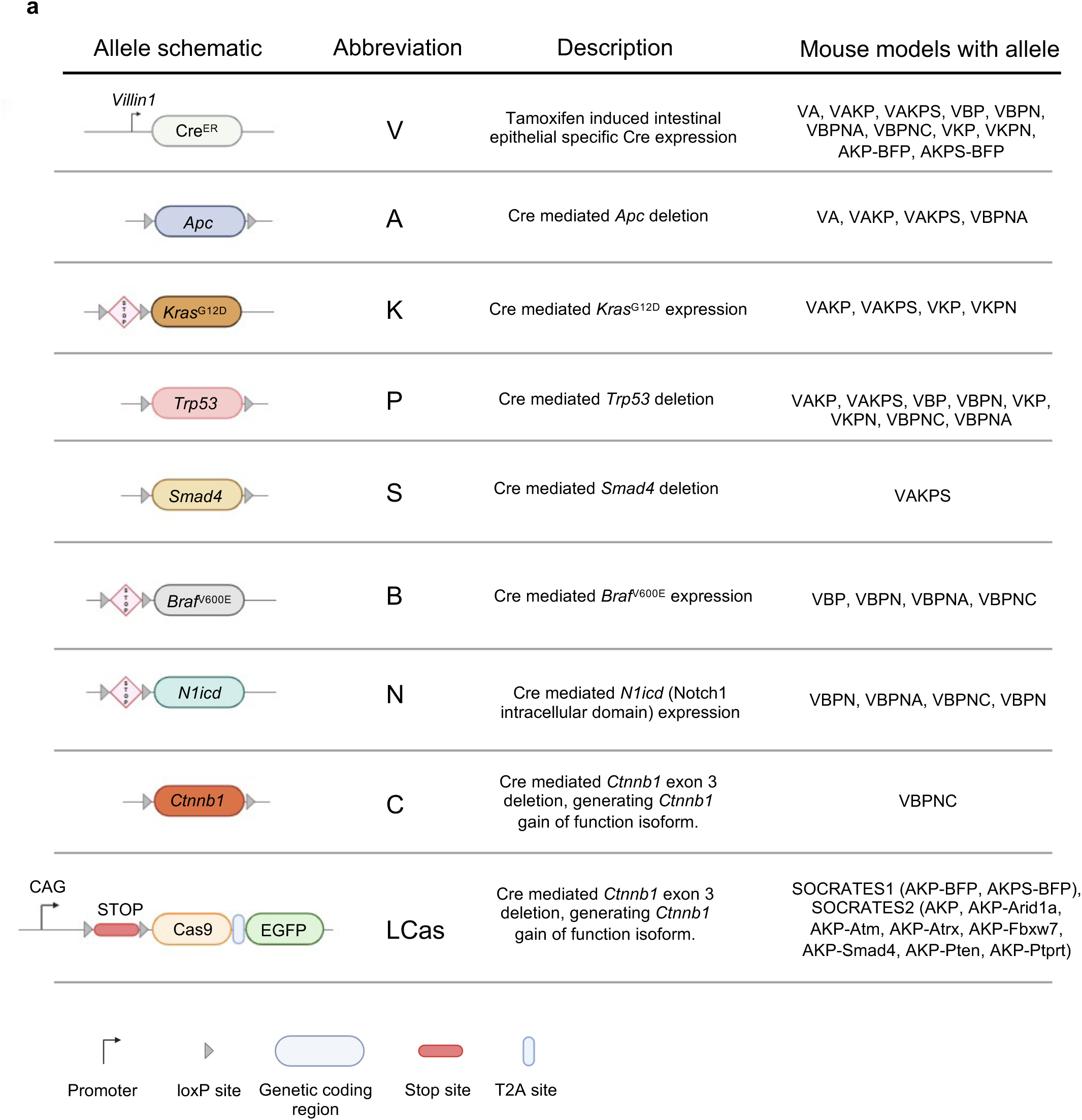
Genetics and transgenes. Table describing the used mouse alleles including abbreviations (EGFP=enhanced green fluorescent protein).

**Extended Data Figure 2.**
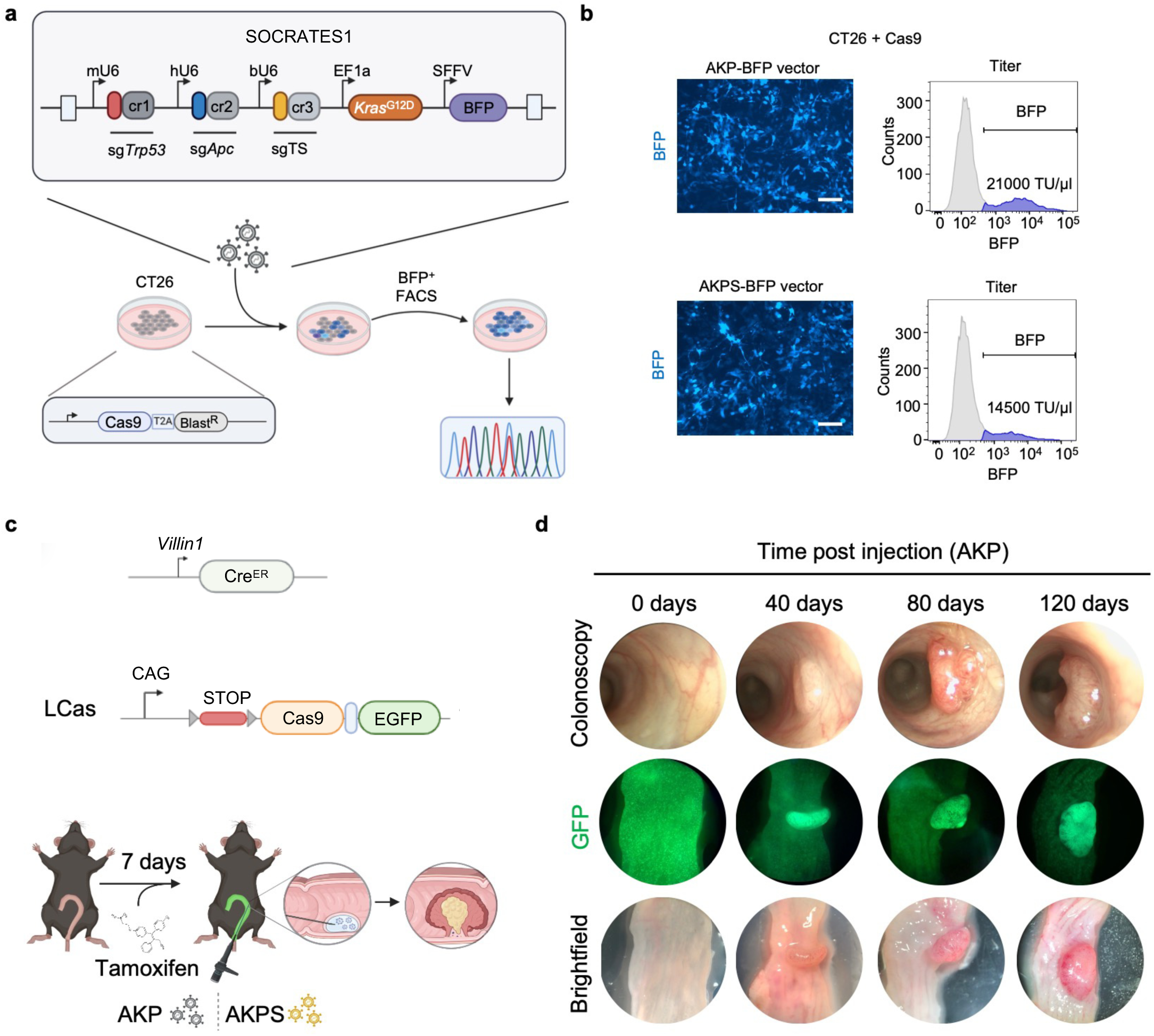
SOCRATES1 vector system development and testing. **a**, Schematic showing the SOCRATES1 vector and depicts the titre testing strategy for this system (single guide RNA of tumour suppressor=sgTS; constant region= cr; blue fluorescent protein=BFP; blasticidin resistance=BLast^R^; fluorescence associated cell sorting=FACS). **b,** Immunofluorescence pictures and flow cytometric analysis of BFP expression testing the *Apc Kras Trp53* (AKP) or *Apc Kras Trp53 Smad4* (AKPS) vectors in CT26 cells. Scale bar 200 µm. **c**, Schematic describing the induction regimen of systemic tamoxifen injection followed by intra colonic guided needle injection of the SOCRATES1 viruses AKP or AKPS. **d,** Colonoscopy, GFP and brightfield images of SOCRATES1 generated tumours with AKP genotype over time.

**Extended Data Figure 3.**
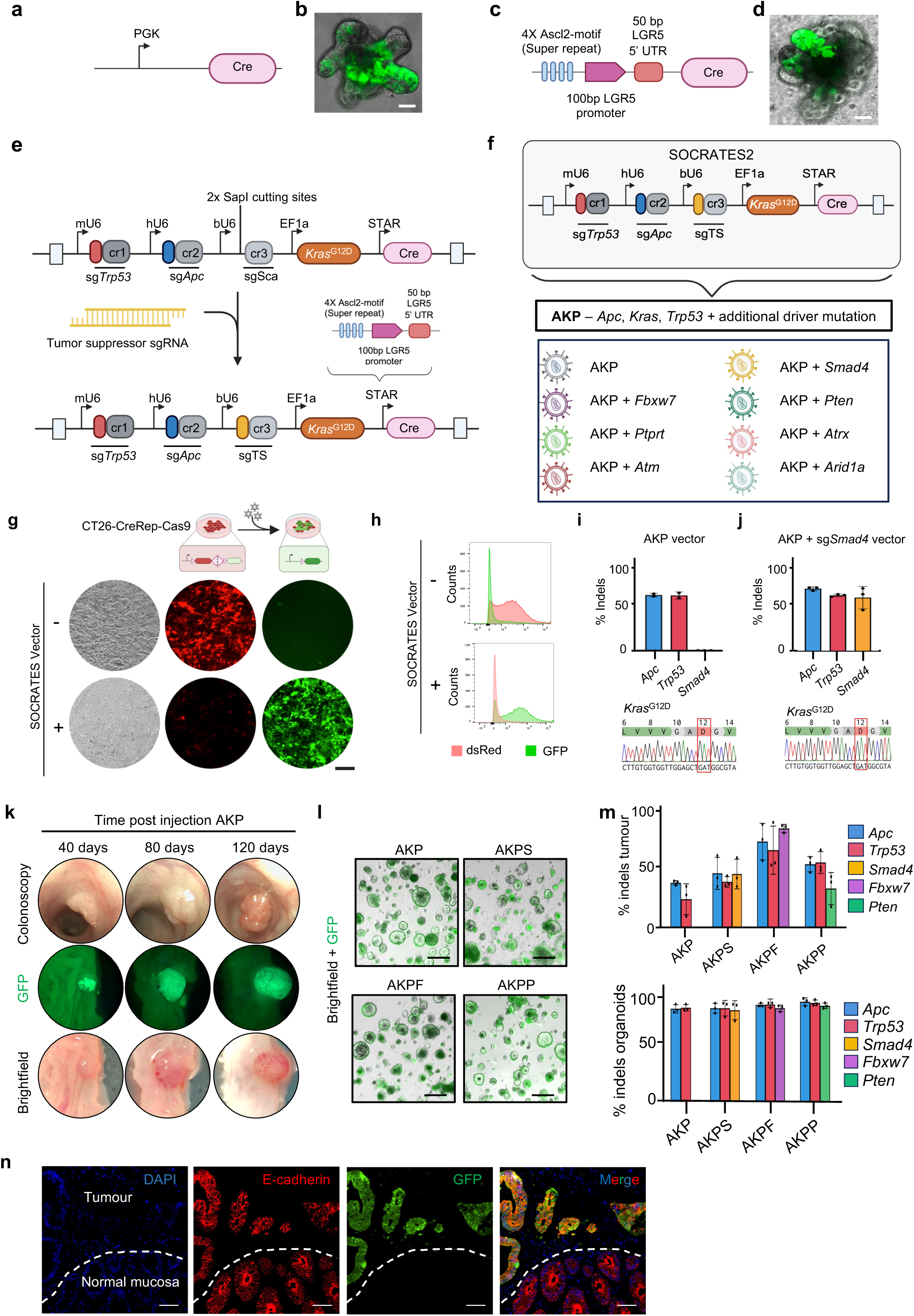
Characterization of SOCRATES2 *in vitro* and *in vivo*. **a**, Cre expression system with PGK promoter. **b,** Microscopy image of mouse small intestinal organoid with LSL-Cas9-GFP transgene after recombination with the construct of a. **c,** STAR promoter and Cre expression system. **d,** Microscopy image of mouse small intestinal organoid with LSL-Cas9-GFP transgene after recombination with the construct of c. **e,** Schematic describing the one-step-cloning strategy for SOCRATES2 (Tumour suppressor sgRNA=sgTS; constant region=cr; sgRNA scaffold=sgSca). **f,** SOCRATES2 vector for modelling advanced cancer with multiplexed and flexible editing option. **g,** Immunofluorescence pictures of CT26-CreRep-Cas9 (tdTomato-LSL-GFP) infected with lentivirus from AKP vector. **h,** Representative histograms of CT26-CreRep-Cas9 cells from g post infection and induction of GFP expression. **i**, Quantification of indel frequency in organoids infected with an AKP construct. (n=2) and chromatogram of KrasG12D mutation. **j**, Quantification of indel frequency in organoids infected with an AKPS construct. (n=3) and chromatogram of KrasG12D mutation. **k,** Colonoscopy, GFP and brightfield images of SOCRATES2 generated tumours with AKP genotype over time. **l,** Immunofluorescence images of organoids derived from SOCRATES2 generated tumours with AKP, AKPS, AKPF and AKPP genotypes. Scale bar 100 µm. **m,** Analysis of SOCRATES mediated genome editing from tumours with indicated genotypes. **N,** Immunofluorescence of AKPS tumour. Dashed line demarks normal tissue. Scale bar 100 µm.

**Extended Data Figure 4.**
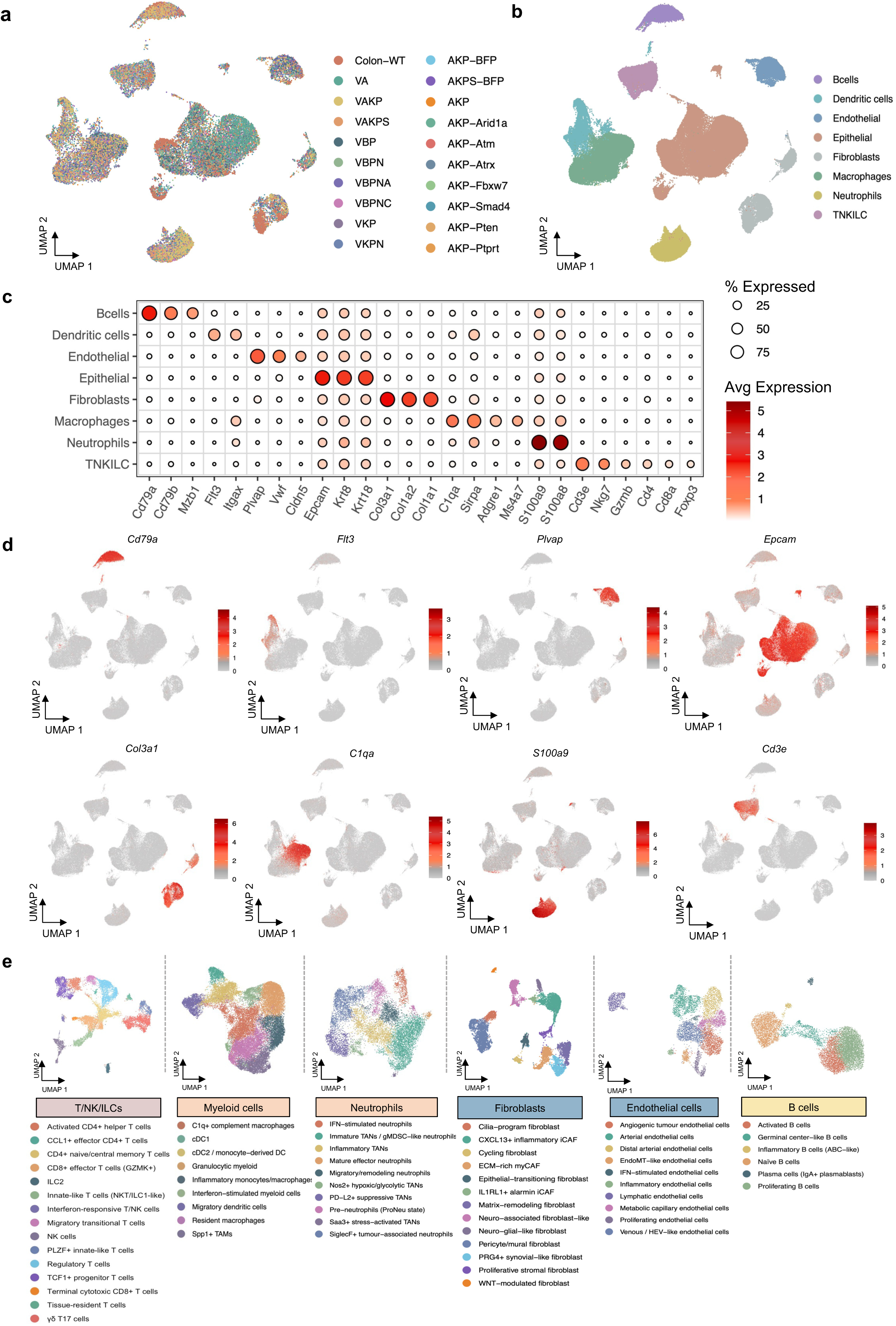
Cell type annotation. **a,** UMAP plot showing individual tumours from the models (131,716 cells). **b and c,** UMAP and bubble plot presentation of marker genes for cell type identification. **d,** UMAPs with marker gene representation. **e,** Cell state polarization of TME cells in the mouse atlas (T/NK cell n=11,548; Myeloid (Macrophages/Dendritic) cells n=29,226; Neutrophils n=11,616; Fibroblasts n=9,965; Endothelial cells n=7,167; B cells n=7,167).

**Extended Data Figure 5.**
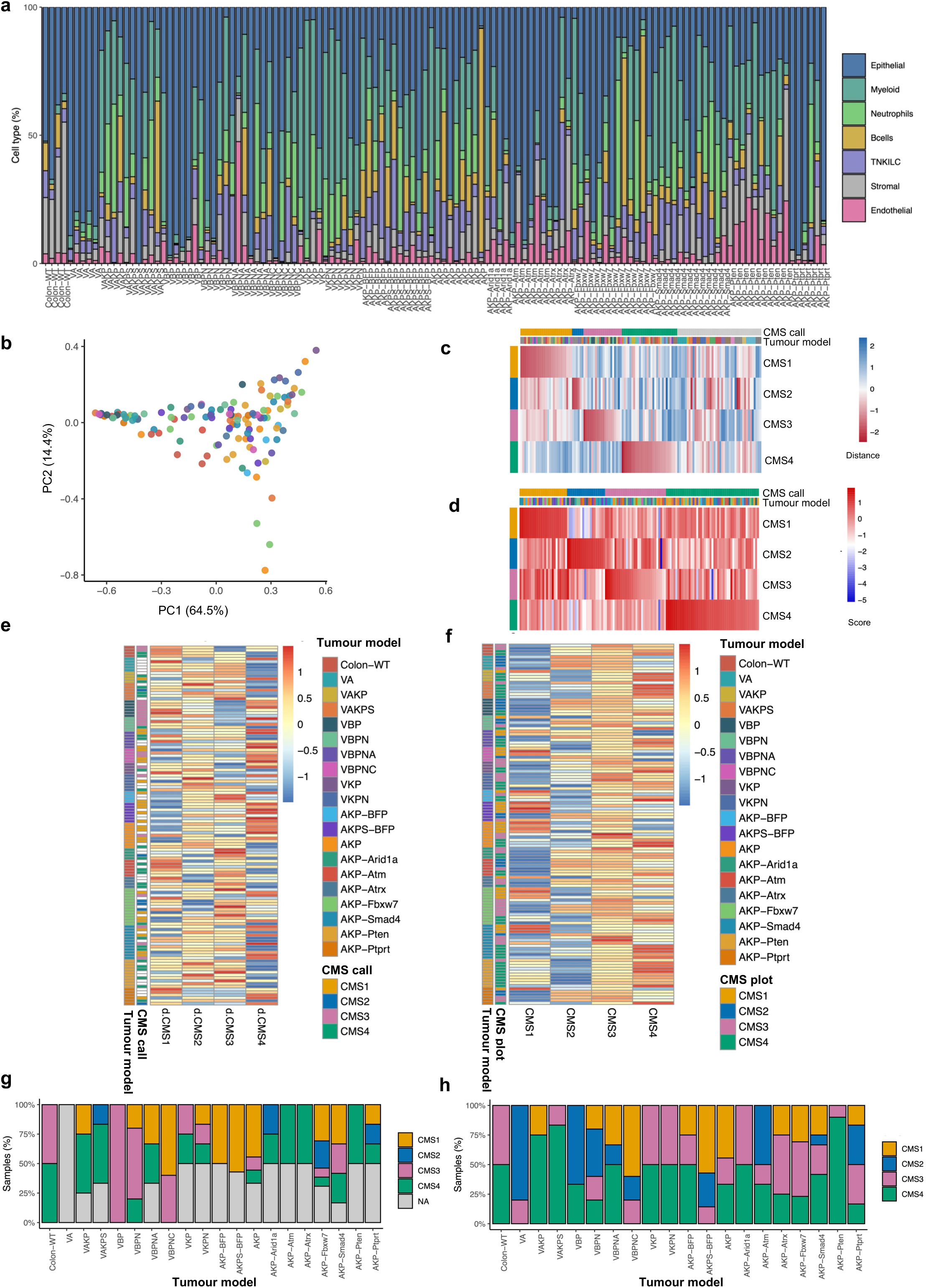
Cell type heterogeneity and consensus molecular subtypes. **a,** Bar plot showing cell type distribution of individual samples (n=126). **b,** PCA plot based on cell type composition of each individual sample. **c,** CMS annotation for pseudo-bulk of each sample using MmCMS classifier. **d,** CMS-like subtypes were inferred by ssGSEA of curated CMS gene-sets on pseudobulk expression profiles, with samples assigned to the highest-scoring program. **e,** Heatmap showing the distribution of CMS assigned tumours with MmCMS classifier. **f,** Heatmap showing the distribution of CMS assigned tumours with ssGSEA of curated CMS gene-sets. **g,** Stacked bar plot depicting the proportion of tumours assigned to each CMS-like program within tumour models based on MmCMS classifier. **h,** Stacked bar plot depicting the proportion of tumours assigned to each CMS-like program within tumour models based on ssGSEA-derived classification.

**Extended Data Figure 6.**
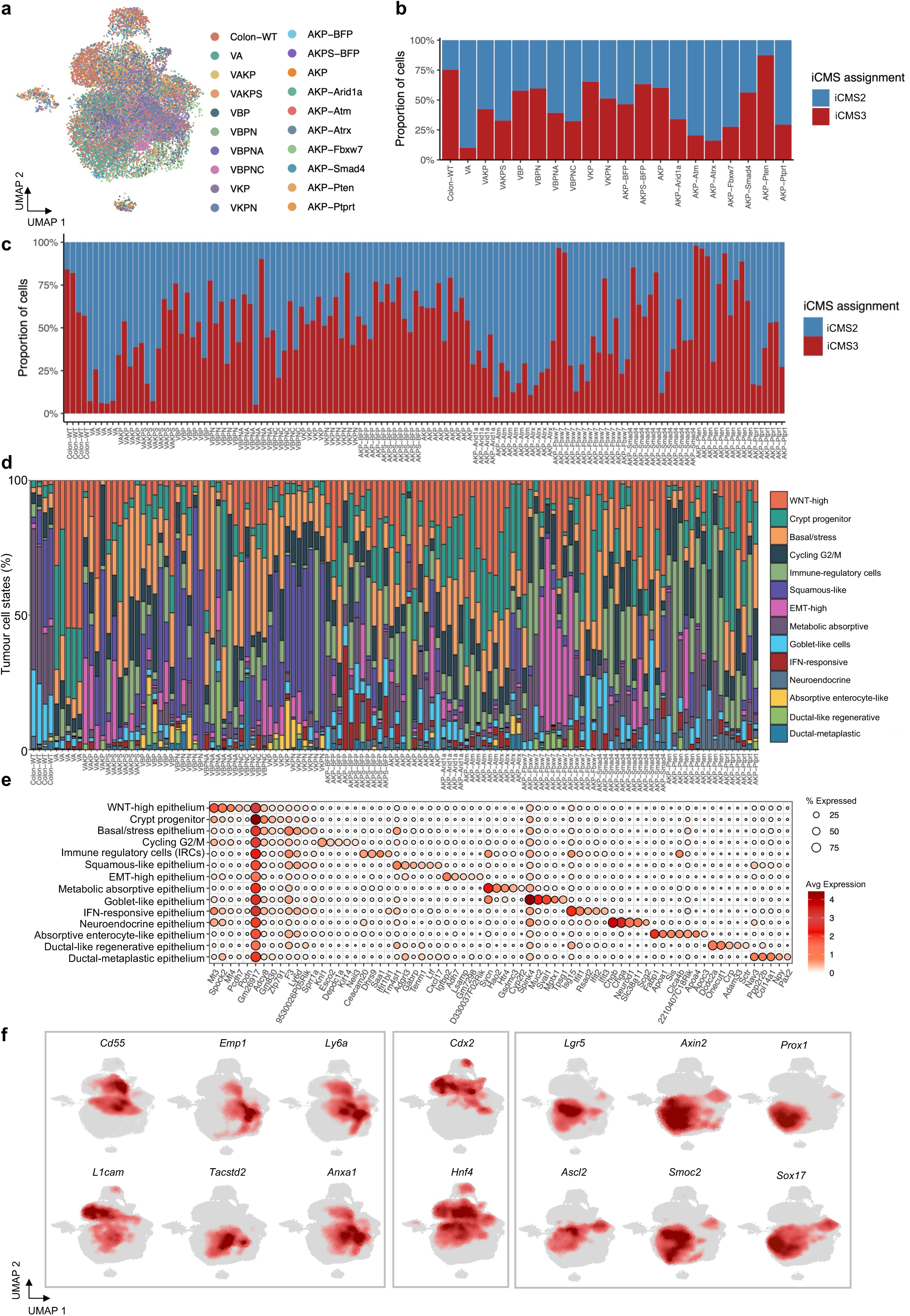
Epithelial cancer cell heterogeneity. **a,** UMAP plot showing individual tumours from the models in the epithelial compartment. **b and c,** iCMS calling per genotype and individual tumour. **d,** Bar plot showing epithelial cell state distribution of individual tumours (n=126). **e,** Bubble plot presentation of marker genes for epithelial cell type identification. **f,** UMAP plots showing epithelial cell state clusters markers.

**Extended Data Figure 7.**
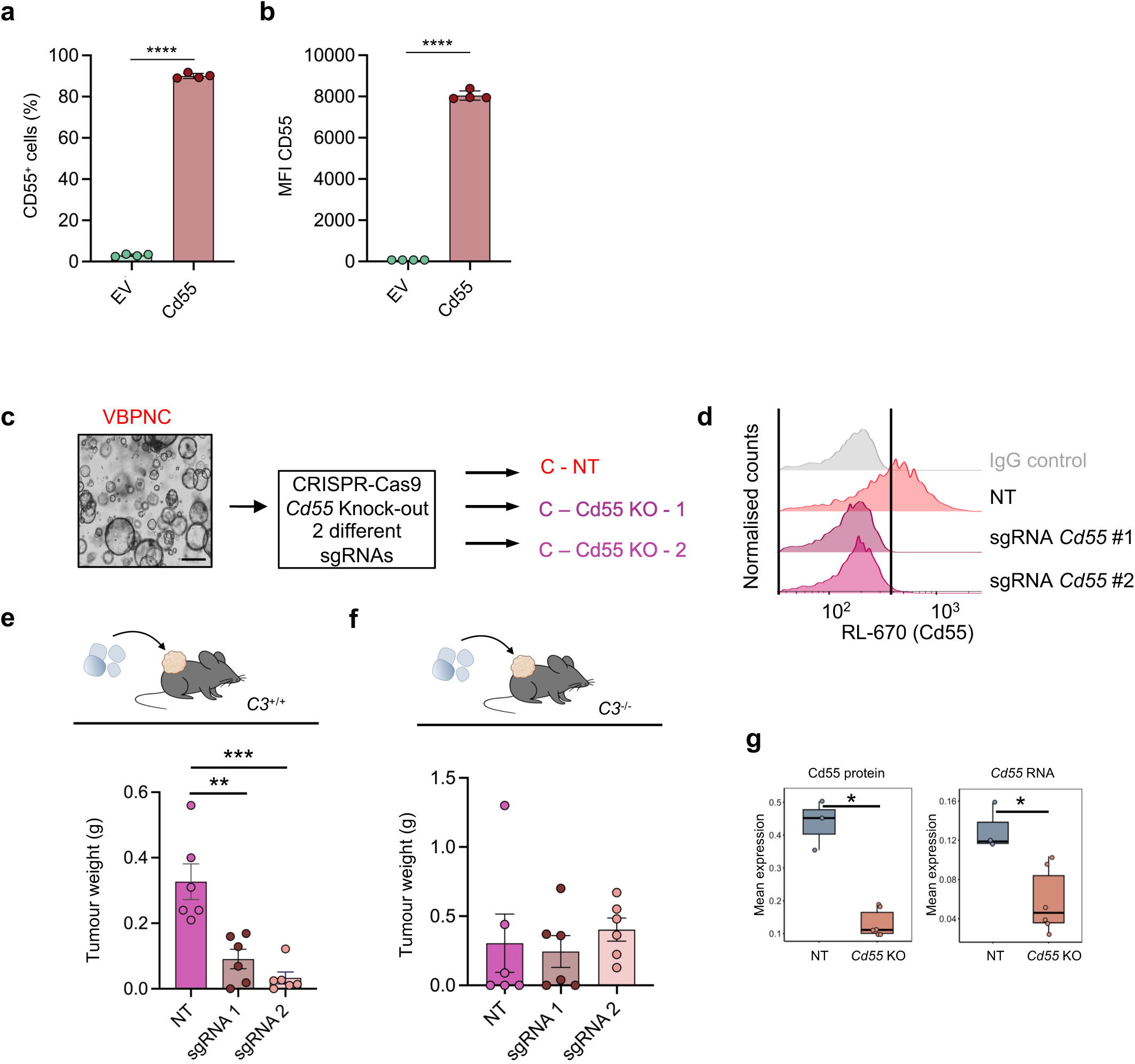
CD55 effect on TME remodelling. **a, b** Flow cytometric quantification of CD55 expression in VBPN EV and VBPN CD55 organoids, shown as the percentage of CD55-positive cells (**a**) and mean fluorescence intensity (MFI) (**b**). **c**, Schematic describing the CRISPR/Cas9 knock-out strategy in VBPNC organoids. **d,** Histogram of *Cd55* expression in non-targeting (NT) sgRNA or *Cd55* sgRNAs. **e,** Tumour weights at endpoint of transplanted organoids from d in C57BL/6 (**e**) and *C3*^-/-^ (**f**) mice (n=6). **g,** Box-plot showing the CITE-seq data for Cd55 protein and *Cd55* mRNA in VBPNA non-targeting (NT) and VBPNA *Cd55* knock out (KO) tumours (n=6). Statistical significance in a, b, e, f and g was assessed using a two-sided unpaired Student’s *t*-test.

## Material and Methods

### Plasmids and molecular cloning

SOCRATES vectors were generated with classical restriction cloning. To avoid recombination during lentiviral packaging and integration due to repetitive DNA sequences we used human, mouse and bovine U6 promoters and three different sgRNA constant regions from pMJ117 (Addgene, #85997), pMJ179 (Addgene, #85996), pMJ114 (Addgene, #85995), respectively. Further, we introduced Cre-recombinase, PGK promoter and Lenti-backbone which were cloned from Lenti-sgNT3/Cre (Addgene, #89654). The STAR promoter with 2x Ascl2 motive was a kind gift from Hugo J. Snippert. This sequence was changed to the STAR promoter with 4 Ascl2 motives to increase expression levels. Kras-G12D sequence was cloned from pLVX-Tight-Puro-KrasG12D2B (Addgene, #139786) and set under the control of a EF1α-core promoter taken from Lenti-Cas9-2A-Bast (Addgene #73310). SFFV-EBFP2 marker cassette for VLCas mice experiments was cloned from LeGO-EBFP2 (Addgene, #85213). NT and sgRNAs targeting *Cd55* were cloned via classical restriction enzyme cloning into pSpCas9(BB)-2A-GFP (PX458) (Addgene #48138).

CD55 overexpression vectors were designed and acquired through VectorBuilder. The murine *Cd55* coding sequence (NM_010016.5) was expressed from the EF1α promoter in the pLV[Exp]-Bsd-EF1A backbone. The corresponding empty vector (pLV[Exp]-Bsd-EF1A>ORF_Stuffer) served as control.

### Virus production

Lenti-X™ 293T cells (Takara Bio, 632180) were cultured in DMEM GlutaMax (Thermo Fisher Scientific, #31966021) with 10% FBS (Thermo Fisher Scientific, #A5256701) and 1% penicillin-streptomycin (Sigma, #P4458-100ML) up to a confluency of 70%.

Lenti-X cells were transfected using polyethyleneimine (Polyscience, #23966-1) with plasmid of interest, pMD2.G and psPAX2 plasmids in a molar ratio of 1:1:1. Supernatant was changed 12-24 hours post transfection. Supernatant was centrifuged at 25,000 rpm, 4°C for 2 hours in an Ultracentrifuge (Beckman Coulter, Optima L-90K) using a SW32TI rotor (Beckmann Coulter). Afterwards supernatant was removed, lentiviral pellets resuspended in ice cold PBS and aliquots were stored at -80°C.

### Cell line generation

CT26 cells were cultured in DMEM GlutaMax (Thermo Fisher Scientific, #31966021) with 10% FBS (Thermo Fisher Scientific, #A5256701) and 1% penicillin-streptomycin (Sigma, #P4458-100ML). For titre testing CT26 cells were transduced with lentivirus produced with Cre reporter plasmid (Addgene #62732) harbouring a LoxP-dsRed-STOP-LoxP-EGFP-IRES-PuromycinR cassette. The cells were selected using Puromycin (Thermo Fisher Scientific, #A1113803) and sorted for dsRed positive and GFP negative cells, resulting in the CT26-CreRep cell line. For mutational testing CT26-CreRep cells were transduced with lentivirus produced with Lenti-Cas9-2A-Bast (Addgene #73310). The cells were selected using Blasticidin (Thermo Fisher Scientific, #A1113903) resulting in the CT26-CreRep-Cas9 cell line.

### Mutational testing pipeline

500,000 CT26-CreRep-Cas9 cells were resuspended in 100 µl PBS and transduced with 5 µl of SOCRATES virus for 20 minutes. After 20 minutes of incubation at RT the cells were washed with PBS, plated and cultured with DMEM GlutaMax (Thermo Fisher Scientific, #31966021) and 10% FBS (Thermo Fisher Scientific, #A5256701). At indicated time points, GFP positive cells were sorted and genomic DNA was isolated. The loci of the intended CRISPR edits were amplified via PCR with loci specific primers (Extended Data Table 2). The purified PCR products were analysed by Sanger sequencing (Eurofins Genomics). The CRISPR indel efficiency was determined via ICE analysis (Synthego).

### Testing of active lentiviral titre

500,000 CT26-CreRep cells were resuspended in 100 µl PBS and transduced with 5 µl of virus. After 20 minutes of incubation at room temperature the cells were washed with PBS, plated and cultured with DMEM GlutaMax (Thermo Fisher Scientific, #31966021) and 10% FBS (Thermo Fisher Scientific, #A5256701). 24 hours post transduction the media was changed. 7 days post transduction cells were harvested using TrypLE Express (Thermo Fisher Scientific, #12605010), washed and resuspended in PBS (2% FBS). Subsequently, the percentage of GFP-positive or BFP-positive cells was determined via flow cytometry analysis and the active lentiviral titre was calculated. For *in vivo* experiments, we used virus batches with an active lentiviral titre >25,000 TU/µl.

### Generation of mouse organoids

Mouse small intestine organoids were generated based on previously published protocols with minor modifications. Briefly, the first 10 cm of the small intestine was excised, opened longitudinally, and mucus was removed. Tissue was cut into ∼5 mm fragments and washed repeatedly in cold PBS until the supernatant appeared clear. After settling, tissue fragments were incubated in PBS containing 2 mM EDTA (Invitrogen, #AM9260G) for 30 minutes at 4 °C on a rotating platform. The EDTA solution was removed, cold PBS was added, and tissue was vigorously pipetted to release crypts. Crypt-containing supernatants were collected, filtered through a cell strainer, and centrifuged to pellet crypts. The pellet was resuspended in Cultrex BME, reduced growth factor matrix (R&D Systems, #3433-005-01), and 20 µl domes were seeded into 6-well plates. Small intestine organoids were cultured in Advanced DMEM/F12 (Gibco, 12634028) supplemented with 1% L-glutamine (Thermo Fisher Scientific, #25030024), 1% HEPES (Sigma, #H0887-100), 1% penicillin–streptomycin (Sigma, #P4458-100ML), 50% WRN-conditioned medium, 1x B27 (Gibco, #12585010), 1x N2 (Gibco, #17502048), 50 ng ml⁻1 EGF (Peprotech, #AF-100-15), 10 µM Y-27632 (Holzel, #M1817), 0.5 µM A83-01 (Sigma, #SML0788), and 2 µl ml⁻1 Primocin (InvivoGen, #ant-pm-2).

### Tissue dissociation

For tissue dissociation into single cells, fragments were thawed and enzymatically digested with the tumour dissociation kit (Miltenyi, #130-095-929) following manufacturer’s instructions. In brief, 5 ml of DMEM GlutaMax (Thermo Fisher Scientific, #31966021) was supplemented with 500 µl of the enzyme mix and placed in the gentleMACS C Tube (Miltenyi, #130-096-334) together with the tumour fragments. The tubes were placed in the gentleMACS Octo Dissociator with heaters (Miltenyi, #130-096-427) and dissociated using the default programs 37C_m_TDK_1.

### Mouse tumour derived organoids

Tumour derived organoids were generated from intestinal tumours of CRISPR-GEMMs. VBPN, VBPNA and VBPNC organoids were previously described^68^. After tumour dissociation, the cells were filtered through a 100 µm cell strainer (Greiner, #542000). Next, the cells were centrifuged down and the pellet resuspended in Cultrex reduced growth factor basement membrane (BME; R&D Systems, #3433-005-01) and the organoids were grown in Advanced DMEM/F12 (Gibco, #12634028) supplemented with 1% L-Glutamine (Thermo Fisher, #25030024), 1% HEPES (Sigma, #h0887-100), B27 (Gibco, #12585010), N2 (Gibco, #17502048) and 1% penicillin-streptomycin (Sigma, #P4458-100ML). Depending on the mutational background of the organoids, the basic media was additionally supplemented with 50 ng/ml EGF (Preprotech, #AF-100-15-1000), 100 ng/ml of Noggin (Peprotech, #250-38-100) or 100 ng/ml of R-spondin conditioned media (U-Protein Express). Organoids were split every 2-4 days.

### Organoid genome editing

Organoid cultures were genetically engineered to generate *Cd55* knock-out lines using CRISPR-Cas9. Organoids were mechanically fragmented and electroporated with plasmids encoding Cas9-GFP as described above and one of two independent *Cd55*-targeting single-guide RNAs, followed by immediate recovery in complete organoid medium and replating in BME. Twenty-four to forty-eight hours after electroporation, organoids were dissociated and FACS was performed to select for GFP-positive cells. Clonal organoids were subsequently isolated and expanded, and genome editing was validated by PCR amplification of the targeted locus and Sanger sequencing. CRISPR-mediated indel frequencies were quantified using ICE analysis (Synthego); due to clonality, indel rates of 0%, 50%, or 100% were expected. Only organoid lines with confirmed biallelic *Cd55* disruption were used for downstream experiments.

### Animal experiments and ethics

All mouse experiments were approved by the local authorities of the Regierungspräsidium Karlsruhe, Baden-Wurttemberg, Germany under the permit numbers G-148-20, G-235/20, G-164/22. Mice were housed according to the local and latest standards at the DKFZ animal facilities with a 12-hour dark and light cycle, a constant temperature (20-24°C) and humidity (45-65%) and were provided with a rodent-specific diet and water *ad libitum*. LCas (B6;129-Gt(ROSA)26Sortm1(CAG-cas9*,-EGFP)Fezh; RRID:IMSR_JAX:024857); VLcas (vilinCreER (B6- Tg(Vil1-cre/ERT2)23Syr; RRID:IMSR_JAX:020282; (B6;129-Gt(ROSA)26Sortm1(CAG-cas9*,-EGFP)Fezh; RRID:IMSR_JAX:024857)); VAKP (B6-Tg(Vil1-cre/ERT2)23Syr Apctm2Rak Krastm4Tyj Trp53tm1Brn); VAKPS (B6-Tg(Vil1-cre/ERT2)23Syr Apctm2Rak Krastm4Tyj Trp53tm1Brn Smad4tm2.1Cxd); VKPN (B6-Tg(Vil1-cre/ERT2)23Syr Krastm4Tyj Trp53tm1Brn Gt(ROSA)26Sortm1(Notch1)Dam; previously described^7^); VKP (B6-Tg(Vil1-cre/ERT2)23Syr, Krastm4Tyj Trp53tm1Brn; previously described^7^); VBPN (B6-Tg(Vil1-cre/ERT2)23Syr Trp53tm1Brn Gt(ROSA)26Sortm1(Notch1)Dam; previously described^68^); VBP (B6-Tg(Vil1-cre/ERT2)23Syr Trp53tm1Brn; previously described^68^); VBPNA (B6-Tg(Vil1-cre/ERT2)23Syr Trp53tm1Brn Gt(ROSA)26Sortm1(Notch1)Dam Apctm2Rak; previously described^68^); VBPNC (B6-Tg(Vil1-cre/ERT2)23Syr Trp53tm1Brn Gt(ROSA)26Sortm1(Notch1)Dam Ctnnb1tm1Mmt; previously described^68^). All the GEMMs were bred on a C57BL/6J background. *villin1*CreER, *Ctnnb1*, *Braf* and *Kras* alleles were kept in heterozygosity, *Trp53* and *Smad4* alleles were kept in homozygosity, whereas the *Apc*^fl^*, Rosa26-*LSL-N1icd *and Rosa26-*LSL-Cas9 allele was utilized in either homozygosity or heterozygosity.

### Colonoscopy-guided submucosal injection of virus or 4-hydroxytamoxifen

Tested virus with a concentration of >25,000 TU/µl was thawed on ice and resuspended. Murine colonoscopy and liquid injection were conducted as previously described^31^. In short, colonoscopy was done using a portable endoscope video unit (Karl Storz, Tele Pack Vet x Led) and a telescope (Hopkins, #67030 BA). Mice were anaesthetized with isoflurane and lentivirus was injected using a syringe (Hamilyon, #7656-01), a transfer needle (Hamilton, 7770-02) and an injection needle (Hamilton, #7803-05). Three bubble-forming injections of 70 µl 4-hydroxy tamoxifen (100 µM) or virus were performed inside the colonic submucosa. Tumour initiation and growth was monitored routinely after the injection, via colonoscopy.

### Subcutaneous injection of tumour-derived organoids

Organoids were dissociated into single cells and resuspended in 100 µl of a 1:1 mixture of PBS and BME. Mice were anesthetized and the single-cell organoid suspension was injected into the left flank using a syringe. Monitoring of the tumour growth was performed twice per week with a tumour measurement minimum once per week.

### Intra-splenic injection of organoids

Organoids were dissociated into single cells and injected in the spleen of the mice using a fine-gauge syringe in a final volume of 50 µl PBS. Access to the spleen was achieved through a small surgical incision in the left lateral abdomen of the mouse, followed by gentle exteriorization of the spleen. 10 mg/kg lidocaine and 3 mg/kg bupivacaine were subcutaneously administrated after surgery close to the incision site. Metamizol was administered in the drinking water for three days. Animals were euthanized four weeks after implantation of the modified tumour organoids, marking the experimental endpoint.

### Genome editing validation

Genomic DNA was isolated from tumour tissue and organoid cultures using the DNeasy Blood & Tissue Kit (Qiagen, #69506) according to the manufacturer’s instructions. Targeted genomic loci corresponding to CRISPR/Cas9 target sites were amplified by PCR using locus-specific primers (Extended Data Table 2). PCR products were purified and subjected to Sanger sequencing (Eurofins Genomics). Genome editing efficiency and the spectrum of CRISPR-Cas9-induced insertions and deletions were quantified from Sanger sequencing traces using the ICE analysis tool (Synthego), enabling determination of indel frequencies at each targeted locus.

### Tissue processing and histopathological analysis

Tissues were fixed in 10% formalin at 4°C overnight. Paraffin-embedded samples were sectioned at 3-5 μm thickness. Before staining, sections underwent deparaffinization through two consecutive xylene washes (3 minutes each), followed by rehydration using a graded ethanol series (100%, 95%, 80%, and 70%, each for 1 minute).

### Immunofluorescence

Following tissue processing, antigen retrieval was performed by steaming the slides in boiling Target Retrieval Solution (Dako, #S1699) for 30 minutes. Afterwards, the slides were cooled for 20 minutes at room temperature, rinsed in distilled water and washed in PBST 0.1% for 5’. Next, sections were blocked in TNB buffer (0.1 M Tris-HCl, pH 7.5, and 0.15 M NaCl containing 0.5% w/v blocking reagent, Perkin Elmer #FP1020) for 1–2 hours at room temperature. For immunostaining, slides were incubated for 1 hour at room temperature or overnight at 4°C with primary antibodies (Extended Data table 4). Following incubation, sections were washed in PBST 0.1% and then incubated with the appropriate secondary antibody (Extended Data table 4) for 30 minutes at room temperature, washed with PBST 0.1% and incubated with DAPI (1:1000) for 10 minutes at room temperature. The slides were washed again in PBST 0.1% and mounted with Fluoromount-G (Southern Biotech #0100-01). Images were acquired using a Zeiss LSM 710 microscope and analysed with FIJI (ImageJ) or QuPath-0.4.3

### *ln situ* hybridization

*ln situ* hybridization (*ISH*) was carried out using the manual RNAscope 2.5 HD Reagent Kit BROWN (Advanced Cell Diagnostics, Hayward, CA, #322300), strictly following the manufacturer’s protocol. Formalin-fixed paraffin-embedded (FFPE) tissue sections (3-5 μm thick) were cut and incubated in a 60°C oven for two hours before staining. To ensure RNA integrity and sample quality, each tissue was first tested with a positive control probe before evaluating experimental results. Additionally, a negative control probe (DapB, Advanced Cell Diagnostics, Hayward, CA, #0043) was included to confirm that observed staining resulted from specific target probe binding rather than non-specific interactions. The probes *Axin2* (#400331) along with the positive control probe *Ppib* (#313911) (both from Advanced Cell Diagnostics, Hayward, CA).

### Immunohistochemistry (IHC)

Following tissue processing, antigen retrieval was performed by steaming the slides in boiling Target Retrieval Solution (Dako, #S1699), or Target Retrieval Solution pH9 (S2367) for 30 minutes. The slides were then cooled for 20’ at room temperature, washed in TBST 0.1% and subsequently incubated in 3% hydrogen peroxide (Sigma, #95321) for 10’. Tissue sections were then incubated in Normal Swine Serum (Biozol, #LIN-ENP9010-10), at room temperature for 1 hour. Primary antibodies (Extended Data table 4) were applied either overnight at 4°C or for 1 hour at room temperature. After incubation, slides were rinsed with TBST 0.1% and incubated for 30 minutes to 1 hour at room temperature together with the appropriate biotin-conjugated secondary antibody (Extended Data table 4). Then, the slides were washed with TBST 0.1% and incubated with the VECTASTAIN® ABC kit (Vector Laboratories, #PK-6100) for 30 minutes at room temperature. The slides were washed again with TBST 0.1%. Signal detection was conducted with DAB chromogen (Agilent Dako, #GV82511-2), following the manufacturer’s instructions.

### Quantitative real-time PCR

Total RNA was isolated from organoid cultures or other cell populations using the RNeasy Mini Kit (Qiagen, 74106) or according to the manufacturers’ instructions. Complementary DNA (cDNA) was synthesized from up to 1 µg of total RNA using the High-Capacity cDNA Reverse Transcription Kit (Applied Biosystems, #4374966). Reverse transcription was performed with incubation at 25 °C for 10 min, followed by extension at 37-42 °C for 60-120 minutes and enzyme inactivation at 85 °C. qPCR was performed using PowerUp SYBR Green Master Mix (Thermo Fisher Scientific, #A25742). Gene expression was normalized to housekeeping genes (s18 or Gapdh for murine samples). Relative expression levels were calculated using the ΔΔCt method and are reported as fold change relative to the indicated control conditions. qPCR oligonucleotide sequences used in this study are shown in Extended Data Table 2.

### Complement deposition assay

Tumour cells were dissociated into single cells and incubated in Advanced DMEM/F12 supplemented with 25% of Pooled Human Complement Serum (Innovative Research, ICSER25ML) for 2-3h at 37°C. As a negative control, serum was heat-inactivated (HIS) at 56 °C for 30 minutes to abolish complement activity. Cells were then washed, and cell-surface C3b deposition was quantified by flow cytometry using the indicated antibodies (Extended Data Table 4).

### Complement-dependent cytotoxicity (CDC)

Tumour cells were dissociated into single cells and incubated with Advanced DMEM/F12 supplemented with 25% of Pooled Human Complement Serum (Innovative Research, ICSER25ML) for 2-3 hours at 37°C. Triton X-100 (2%) was added during the final 15 min as a 100% lysis control. Heat-inactivated serum (HIS) was included as a negative control. Cell lysis was measured using the ToxiLight^®^ Non-Destructive Cytotoxicity BioAssay Kit (Lonza, LT17-217). CDC was calculated as:

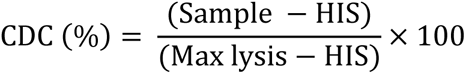

### Single-cell RNA sequencing sample preparation

Tumour samples were dissociated into single cells as previously described. Samples were multiplexed using a maximum of eight different hashing antibodies (TotalSeq^TM^-B, BioLegend) according to the manufacturer’s protocol. In brief, single cells were resuspended in PBS, 2% FBS (Thermo Fisher Scientific, A5256701), blocked with 1:50 FcR Receptor Blocking Reagent Mouse (Miltenyi Biotec, #130-09 2-575) and incubated for 10 minutes on ice. Next the antibodies were added at the recommended concentration and incubated at 4°C for 30 minutes. Afterwards, the cells were washed two times with ice cold PBS + 4% FBS. Cells were stained with ZombieNIR (Biolegend, #423105) in ice cold PBS + 2% FBS. Samples were sorted for live cells (ZombieNIR negative). Multiplexed samples were sorted in a single tube. Library generation was performed following the Chromium Next GEM Single Cell 3’ Reagent Kit v3.1 or v4 (Dual Index) protocol, resulting in a library for gene expression (GEX) and a library for hashtag oligos (HTO) per chemistry.

### FACS

CT26, CT26-Rep, CT26-Rep-Cas9, VBPNA NT, VBPNA sgRNA *Cd55* #1, VBPNA sgRNA *Cd55* #2, VBPNC NT, VBPNC sgRNA *Cd55* #1, VBPNC sgRNA *Cd55* #2, tumour cells were prepared as described above. Cells were sorted with FACS Aria I, II and III, FACS Aria Fusion (BD) and analysed with LSRII, LSR Fortessa. The softwares used were BD FACSDiva v8.0.3 and FlowJow V 10.5.3. For gating of CT26 the following gating strategy was applied: FSC-A vs. SSC-A: Cells, SSC-A vs. SSC-W: Single cells, FSC-A vs. BL-525/50: GFP+/- cells, FSC-A vs. YG-586/15: dsRed +/-cells, FSC-A vs. VL-450/50: BFP+/- cells. For gating of VBPNA and VBPNC cells the following gating strategy was applied: FSC-A vs. SSC-A: Cells, SSC-A vs. SSC-W: Single cells, FSC-A vs. RL-780/60: life cells, BL-525/50: GFP +/- cells. For gating of dissociated primary tumour cells the following gating strategy was applied: FSC-A vs. SSC-A: Cells, SSC-A vs. SSC-W: Single cells, FSC-A vs. RL-780/60: life cells. Cells were sorted in FACS buffer for further processing.

### CITE-seq with cell hashing sample preparation

For simultaneous transcriptomic, surface protein, and sample identity profiling, tumour-derived single-cell suspensions were prepared as described above. Following dissociation, cells were subjected to antibody-based cell hashing using TotalSeq™-B hashing antibodies (BioLegend) according to the manufacturer’s protocol. Briefly, cells were Fc receptor–blocked, incubated with hashing antibodies for 30 minutes at 4 °C, and washed thoroughly in ice-cold FACS buffer. Hashed samples were then pooled and viable cells were identified by ZombieNIR (Biolegend, #423105) exclusion and sorted into a single tube. Subsequently, a minimum of 500,000 pooled viable cells were incubated with a CITE-seq antibody cocktail (TotalSeq™-B Mouse Universal Cocktail v1.0; BioLegend, #199902) for 30 minutes at 4 °C, followed by three washes in FACS buffer and resuspension in PBS supplemented with 2% FBS. CITE-seq libraries were generated using the Chromium Next GEM Single Cell 3’ Reagent Kit v3.1 or v4 (Dual Index; 10x Genomics), yielding one library for gene expression (GEX) and one combined library for antibody-derived tags and hashtag oligonucleotides (ADT/HTO) per sample.

### Single-cell RNA sequencing

Single-cell libraries were sequenced with paired-end, dual indexing on a NovaSeq6000 with the following number of cycles: 28 cycles read 1, 10 cycles i7 Index, 10 cycles i5 index and 90 cycles Read 2. It was aimed for a sequencing depth of ∼30,000 read pairs per cell and ∼500 read pairs per antibody.

### Aligning and demultiplexing single-cell RNA-seq reads

Sequencing reads were mapped to the respective reference genome (mouse genome assemblies refdata-gex-mm10-2020-A, respectively) using 10X Genomics Cell Ranger (v6.1.0). The samples were demultiplexed on the feature barcode sequence assigned to the samples via the TotalSeq^TM^-B hashing antibodies in the sample preparation.

### Pre-processing and quality control of single cell data

Sample processing and analysis were performed in R using Seurat. For each sequencing run, Cell Ranger “filtered_feature_bc_matrix” outputs were imported and a Seurat object was created from the Gene Expression matrix. Quality control was performed at the run level by computing standard metrics, including the fraction of mitochondrial (percent.mt; genes matching Amt-) and ribosomal transcripts (percent.ribo; genes matching ARpllARps), and filtering cells based on detected features (nFeature_RNA) and mitochondrial content to remove low-quality profiles and outliers. When available, hashtag oligonucleotide (HTO) counts from the Antibody Capture matrix were added as a separate assay, normalized by centred log-ratio (CLR), and demultiplexed with HTODemux using a stringent positive-quantile threshold; only HTO-classified singlets were retained. For runs with weak hashing performance, HTO features were pre-screened using abundance- and prevalence-based summaries (total counts and fraction of positive cells), and low-information hashtags were excluded prior to demultiplexing. HTO singlets were then processed with standard Seurat workflows (log-normalization, variable feature selection by the “vst” method, scaling of variable genes, PCA, UMAP, and graph-based clustering). Doublets were identified per run using scDblFinder after conversion to a SingleCellExperiment object; to reduce overcalling in dense tumour datasets, scDblFinder classifications were optionally “softened” post hoc by re-labelling only the highest-scoring fraction of cells as doublets to match a fixed target rate, and cells labelled as doublets were removed.

### Merging, data integration and clustering and visualization

All runs were merged using merge with run-specific cell ID prefixes, and run provenance (run and, where applicable, GEM/library identifiers) was parsed from cell names and stored in metadata for batch-aware processing. After merging, additional run-aware QC was applied using per-run, distribution-based thresholds on nFeature_RNA, nCount_RNA, and percent.mt (including removal of mitochondrial outliers), followed by batch correction with Harmony using run as the batch variable. The Harmony embedding was used for UMAP visualization, neighbour graph construction, and clustering. Finally, residual low-quality clusters enriched for low complexity or high mitochondrial content were identified by QC feature distributions and removed, and the dataset was reprocessed to generate the final cleaned object for downstream annotation and analysis.

### Cell type and cell state annotation

Cell type and cell state annotations were assigned based on cluster-specific gene expression patterns and curated marker gene and gene set signatures. Differentially expressed genes for each cluster were identified using Seurat’s FindMarkers function, applying a Wilcoxon rank-sum test with Benjamini–Hochberg correction for multiple testing. Clusters were annotated by integrating differential expression results with established lineage and state-specific marker genes (Extended Data Fig. 6). For downstream analyses, annotated clusters were subset into major cell types, and within each subset highly variable genes were re-identified (500–3,000 genes, depending on subset size). Each subset was then independently reprocessed using the same dimensionality reduction, batch correction, and clustering workflow described above to resolve finer-grained cell states. The relative abundance of cell states across tumour samples was quantified and visualized as stacked bar plots. Gene sets used in this study are provided in Extended Data Table 1, and gene set activity scores were computed at single-cell resolution using the UCell package.

### Differential expression analysis and hierarchical clustering

Differential expression analysis was performed via the Seurat package after generating pseudobulks from the single cell data, considering the sample origin and the desired contrast conditions for the grouping. Pseudobulks were generated as a sum of counts using the AggregateExpression function. Further analysis of the counts data from pseudobulk RNA-seq was performed with DESeq2. The Log2 fold changes were calculated and gene set enrichment analysis (GSEA) performed with the fgsea package (gene ranking based on the Log2 fold change). Gene sets were derived from the molecular signatures database (MSigDB) and literature (Extended Data Table 1).

### Pseudotime analysis

Pseudotime trajectory analysis was performed to model transcriptional state transitions within tumour epithelial cells across disease progression using Seurat, SingleCellExperiment, and Slingshot. Tumour models were grouped into three biologically defined disease stages based on genotype and histopathological characteristics (wild-type [WT], adenoma, carcinoma), assigned at the single-cell level via tumour model metadata and encoded as an ordered factor (WT → Adenoma → Carcinoma). For lineage inference, Seurat objects were converted to SingleCellExperiment format and Slingshot was applied using precomputed cell clusters as graph nodes and Harmony-corrected embeddings as input. Root clusters were manually defined based on enrichment for WT-stage cells, thereby constraining trajectories to originate from normal or early epithelial states and progress toward adenoma- and carcinoma-associated programs. Lineage-specific pseudotime values were projected onto UMAP embeddings for visualization, and Slingshot lineage curves were overlaid by re-running Slingshot in UMAP space to obtain smooth trajectory coordinates consistent with the visualization layout. In parallel, pseudotime centerlines were independently computed by binning cells along pseudotime and averaging UMAP coordinates within adaptive bins, providing a robust representation of dominant trajectory paths independent of Slingshot spline fitting. To characterize transcriptional dynamics, gene expression values and module scores were z-score–normalized per gene set module score and analysed along lineage-specific pseudotime using generalized additive models or equal-width pseudotime binning, with mean scaled expression and contributing cell numbers summarized per bin. All pseudotime analyses were performed in a lineage-resolved manner, and expression trends were interpreted within the inferred lineage structure rather than assuming a single linear trajectory.

### CITE-seq data analysis

CITE-seq datasets were processed in Seurat by importing Cell Ranger output matrices and retaining gene expression (GEX/RNA), hashtag oligonucleotide (HTO), and antibody-derived tag (ADT) layers. Run-level objects were merged with unique cell ID prefixes to preserve sample provenance. RNA data were used for cell-level quality control, normalization, dimensionality reduction, graph construction, clustering, and UMAP visualization at the global dataset level. HTO information was used to assign cells to experimental conditions (NT versus CD55 KO). ADT counts were analysed as a separate protein modality to quantify surface marker expression. For selected cell type-specific analyses, both GEX and ADT layers were jointly used for dimensionality reduction, neighbour graph construction, UMAP visualization, clustering, and marker expression analyses, enabling integrated transcriptomic and proteomic characterization of cellular states.

### Publicly available datasets and human data analysis

Publicly available single-cell data from^39^ were downloaded in AnnData format and imported into R using Bioconductor/Seurat-compatible tooling. Expression matrices and accompanying cell-level annotations were converted into a Seurat object while preserving the original study metadata (e.g., patient/sample identifiers, stage/class labels, and hierarchical cell-type annotations). Where available, the published low-dimensional embedding (UMAP) was transferred from the AnnData object to enable consistent visualization of the reference atlas. To compare global shifts in tissue composition across disease classes, cell-type frequencies were summarized per sample state and a stacked bar plot generated. In addition, a joint human-mouse PCA was performed based on a combined composition matrix (sample x cell type proportion) by harmonizing broad cell-type labels between datasets, constructing matched composition matrices for human and mouse samples (sample x cell type proportion), aligning cell-type columns, and running PCA on the combined sample-by-cell-type proportion matrix to enable cross-species comparison of cell-type composition.

### TCGA COAD analysis

Bulk RNA-seq and clinical data from the TCGA colorectal adenocarcinoma (COAD) cohort were obtained using TCGA biolinks and restricted to primary tumour samples. Raw count data were used to assign consensus molecular subtypes (CMS) with the CMScaller framework, and CMS calls were collapsed to one subtype per patient. Gene signature scores were calculated from normalized expression data (logCPM) as the mean expression of predefined gene sets, and differences in signature activity across CMS classes were assessed using non-parametric statistical tests. For survival analyses, patients were stratified into high- and low-expression groups based on the top and bottom 30% of signature scores. Overall survival was evaluated using Kaplan–Meier analysis and log-rank testing, with effect sizes estimated by Cox proportional hazards models. Associations between gene signature scores and clinicopathological features were further evaluated, including correlations with AJCC pathological stage. In addition, consensus molecular subtypes (CMS) were assigned from raw count data using CMScaller, and signature scores were compared across CMS classes using non-parametric statistical tests.

### Visualisation of the data

Data visualization was performed in R using ggplot2 as the primary plotting framework. Single-cell–specific visualizations, including UMAP embeddings, feature expression plots, and cell-type composition displays, were generated using Seurat and SCpubr. Multi-panel figures were assembled using patchwork, and statistical annotations were added with ggpubr. Heatmaps were generated using ComplexHeatmap. Color palettes and scaling were managed using Polychrome, introdataviz, RColorBrewer, and scales to ensure consistent and accessible colour usage across figures. Rasterization of large point clouds was performed with ggrastr, and selected enrichment- and multivariate-style visualizations were generated using fgsea and fmsb, respectively.

### Statistical analysis

Statistical analyses of single-cell RNA-seq data were performed in R as described in the corresponding Methods sections. Differential gene expression and marker analyses were conducted using non-parametric tests, primarily the Wilcoxon rank-sum test, with multiple testing correction applied where indicated. Comparisons of gene expression levels or module scores between two groups were assessed using Wilcoxon rank-sum tests, while comparisons across more than two groups were performed using Kruskal–Wallis tests, followed by post hoc testing when applicable. Correlations between continuous variables were evaluated using Spearman’s rank correlation. For survival analyses in TCGA cohorts, Kaplan–Meier analysis with log-rank testing was used to assess differences in overall survival between groups, and Cox proportional hazards models were applied to estimate hazard ratios. Statistical differences between expression measurements obtained by flow cytometry and qPCR, tumour measurements, and metastatic burden quantification were calculated using GraphPad Prism (version 10; GraphPad Software). Unless stated otherwise, additional statistical analyses were performed using GraphPad Prism (version 10; GraphPad Software). The specific statistical tests applied, number of biological replicates, and exact *P* values are reported in the figure legends. Data are presented as mean ± standard error of the mean (SEM), and statistical significance is indicated directly in the figures.

## Notes

### Competing Interest Statement

The authors have declared no competing interest.

